# Dual-color fluorescent reporters resolve Alu and LINE-1 retrotransposition in single cells

**DOI:** 10.64898/2026.09.26.754642

**Authors:** Shota Amano, Kazuki Ohnishi, Kei Nishimori, Tomoichiro Miyoshi

## Abstract

LINE-1 (L1) is transcribed by RNA polymerase (Pol) II, whereas the non-coding Short INterspersed Element (SINE) Alu is transcribed by Pol III and exploits L1 ORF2p for mobilization. Conventional L1 retrotransposition reporters rely on spliceosome-dependent introns and cannot directly monitor Alu retrotransposition. Here, we developed *EGFP*^Tet^, a fluorescent SINE reporter in which *EGFP* is interrupted by a *Tetrahymena thermophila* group I self-splicing intron. *EGFP*^Tet^ enabled flow cytometry, fluorescence microscopy, cell sorting, and time-course analysis of Alu mobilization without colony formation. Reporter activation required ORF2p reverse transcriptase activity, and recovered Alu-*EGFP*^Tet^ insertions displayed retrotransposition hallmarks. *EGFP*^Tet^ quantified Alu retrotransposition across cell lines and host-factor perturbations and was repurposed to detect mobilization of the phylogenetically distinct mouse B2 SINE. Defined amino acid substitutions generated an mTurquoise2 derivative without redesigning the self-splicing junctions, allowing dual-color analysis of Alu trans-mobilization and L1 cis retrotransposition in individual cells. In this assay, wild-type L1 supported a higher Alu retrotransposition frequency than the ORF1p-mutant L1. Despite their shared dependence on ORF2p, Alu co-expression did not significantly reduce L1 retrotransposition, and Alu/L1 dual-positive counts exceeded independence-based expectations, indicating preferential co-occurrence of the two reporter signals. *EGFP*^Tet^ provides a fluorescent platform for investigating the shared and distinct cellular conditions that support SINE and L1 mobilization.

## Introduction

Transposable elements have profoundly shaped mammalian genome evolution and continue to affect genome integrity. In the human genome, Long INterspersed Element-1 (LINE-1 or L1) is the only autonomous retrotransposon known to remain active (Luqman-Fatah & Miyoshi, 2023; Richardson et al., 2015). L1-derived sequences account for approximately 17% of the genome (Hoyt et al., 2022). A full-length, retrotransposition-competent L1 encodes ORF1p and ORF2p, with ORF2p providing the endonuclease (EN) and reverse transcriptase (RT) activities required for target-site primed reverse transcription (TPRT) (Cost et al., 2002; Feng et al., 1996; Mathias et al., 1991; Moran et al., 1996). L1-encoded proteins can also mobilize non-autonomous retrotransposons, including Alu and SVA, which comprise ∼10% and ∼0.15% of the genome, respectively (Hoyt et al., 2022). Together, L1 and Alu sequences account for ∼27% of the human genome. *De novo* insertions of these elements can disrupt genes, alter splicing, and generate structural variation (Gilbert et al., 2002; Hancks & Kazazian, 2016; Symer et al., 2002). For example, a hominoid-specific Alu insertion in *TBXT* has also been implicated in the evolution of tail loss (Xia et al., 2024). Retrotransposons are therefore a double-edged source of genomic variation, yet the mechanisms controlling their mobilization remain incompletely understood.

Although both L1 and Alu retrotransposition depend on ORF2p, they differ fundamentally in sequence, expression, and host-factor requirements. L1 RNA is a protein-coding RNA polymerase II transcript (Luqman-Fatah & Miyoshi, 2023), while Alu RNA is a short non-coding RNA polymerase III transcript derived from 7SL RNA and exploits ORF2p for mobilization (Dewannieux et al., 2003; Ullu & Tschudi, 1984; Ullu & Weiner, 1985). The encoded 3′ poly(A) tract is a cis-acting sequence required for efficient Alu trans-mobilization and is thought to facilitate ORF2p recruitment (Dewannieux et al., 2003; Dewannieux & Heidmann, 2005b). Previous work provided evidence for competition between L1 and Alu RNAs for ORF2p through their 3′ poly(A) tracts (Doucet et al., 2015). How this shared dependence relates to their retrotransposition frequencies and co-occurrence within individual cells remains unclear. Moreover, recent comparisons between HeLa cell lines showed that L1 retrotransposes in both HeLa-HA and HeLa-JVM cells, whereas Alu retrotransposition is efficient in HeLa-HA but strongly restricted in HeLa-JVM cells (Moldovan et al., 2024). This finding suggests that cellular requirements for Alu mobilization are at least partly distinct from those governing L1 mobilization.

Established L1 reporters use an antisense indicator cassette interrupted by a sense-oriented spliceosome-dependent intron. The indicator cassette becomes functional only after RNA processing, reverse transcription, and genomic insertion (Kannan et al., 2017; Mita et al., 2018; Moran et al., 1996; Morrish et al., 2002; Ostertag et al., 2000; Terasaki et al., 2013; Xie et al., 2011). These reporters have enabled quantitative measurement of L1 retrotransposition (Ostertag et al., 2000; Xie et al., 2011), recovery of retrotransposed cells (Gilbert et al., 2002, 2005; Symer et al., 2002), and genetic screens for host regulators (Liu et al., 2018; Mita et al., 2020). However, the same strategy cannot be applied directly to Alu because Pol III-transcribed Alu RNAs do not undergo canonical spliceosome-dependent pre-mRNA splicing. To address this limitation, the established *neo*^Tet^ Alu assay incorporated a *Tetrahymena thermophila* 26S rRNA group I self-splicing intron within a neomycin-resistance cassette (Dewannieux et al., 2003; Esnault et al., 2002). As group I introns are removed autocatalytically rather than by the spliceosome (Kruger et al., 1982), *neo*^Tet^ is compatible with Pol III-transcribed Alu RNA. The assay has provided important insights into Alu mobilization (Bennett et al., 2008; Dewannieux et al., 2003; Kroutter et al., 2009; Moldovan et al., 2024).

However, *neo*^Tet^ relies on antibiotic selection and colony formation and does not support direct fluorescence-based quantification, cell sorting, or spectral multiplexing. Here, we developed *EGFP*^Tet^, a fluorescence-based SINE retrotransposition reporter containing a *Tetrahymena* group I self-splicing intron within *EGFP*. We show that *EGFP*^Tet^ detects bona fide ORF2p-dependent de novo Alu insertions, supports quantitative analysis of host-factor requirements and temporal reporter activation, and can be adapted to multiple active Alu subfamilies and the mouse B2 SINE. We further generated an mTurquoise2-based derivative for concurrent detection of Alu trans-mobilization and L1 cis retrotransposition in individual cells. Together, these reporters provide a modular platform for direct detection, sorting, temporal analysis, and spectral multiplexing of Pol III-transcribed SINE retrotransposition.

## Results

### Development of an *EGFP*^Tet^ reporter for visualizing Pol III-transcribed SINE retrotransposition

We selected EGFP for the SINE retrotransposition reporter because it supports rapid fluorescence-based quantification, microscopy, and cell sorting, and can be converted into spectrally distinct variants through defined amino acid substitutions for multicolor detection (Heim & Tsien, 1996). To visualize retrotransposition of Pol III-transcribed SINEs such as Alu in living cells, we adapted a group I self-splicing intron derived from the 26S rRNA of *Tetrahymena thermophila* for insertion into the *EGFP* coding sequence (Figure 1a) (Dewannieux et al., 2003; Esnault et al., 2002). Accurate and efficient self-splicing of this intron requires base pairing between the internal guide sequence (IGS) and flanking exon sequences to form the P1 and P10 helices, including an essential G:U wobble base pair at the 5′ splice site (Barfod & Cech, 1989; Suh & Waring, 1990; Waring et al., 1986). We therefore modified the flanking *EGFP* exon sequences and the intron IGS to preserve P1 and P10 base pairing and the essential G:U wobble base pair without altering the encoded *EGFP* amino acid sequence (Figure 1b).

**Figure 1.**
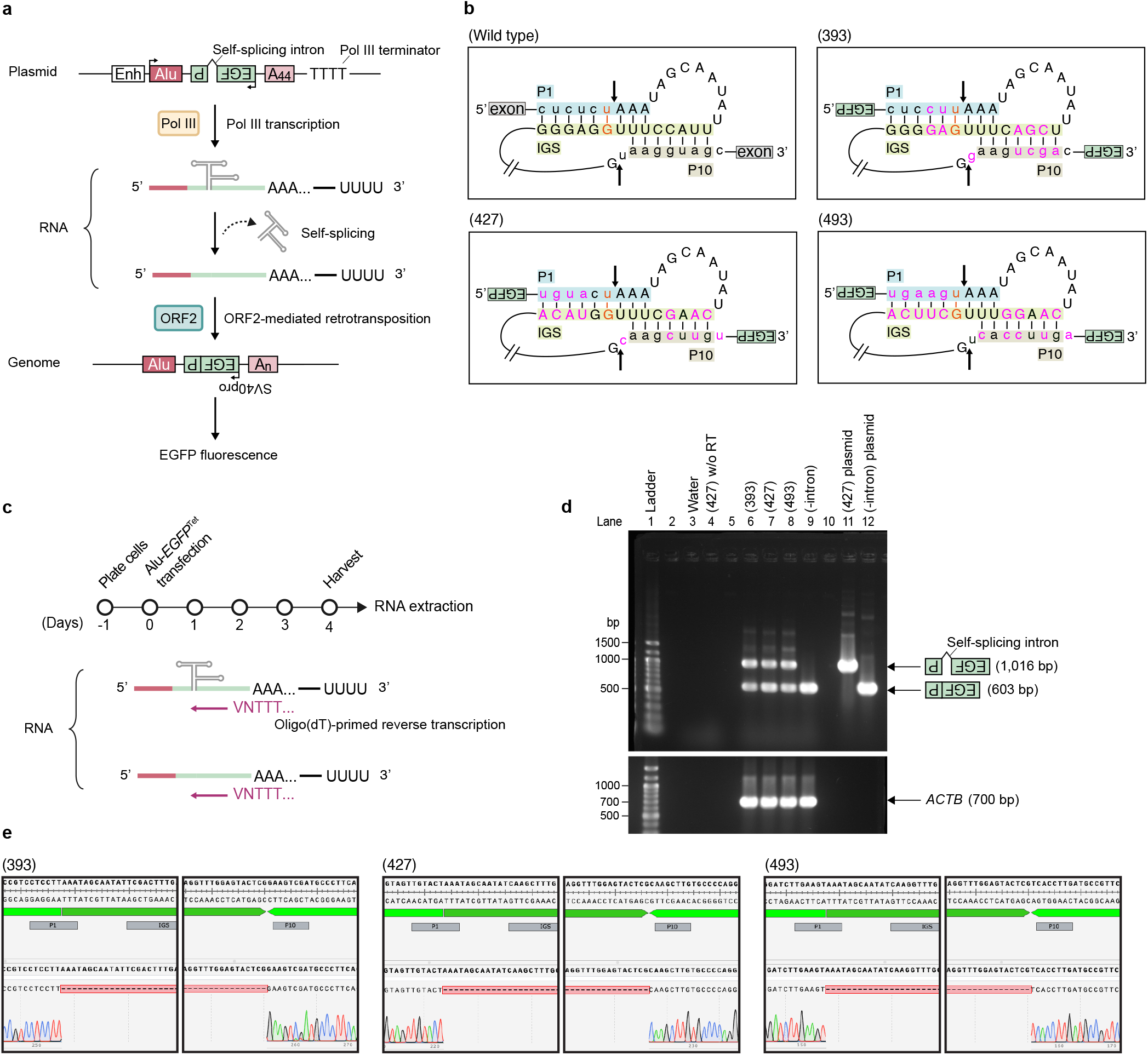
Construction and validation of a SINE-*EGFP*^Tet^ reporter incorporating a *Tetrahymena* self-splicing intron. **(a)** A plasmid map of the Alu-*EGFP*^Tet^ reporter and schematic of reporter activation following L1 ORF2p-mediated retrotransposition. The *EGFP*^Tet^ cassette is placed between the AluY sequence and the encoded A44 tract in the antisense orientation relative to Alu transcription, and *EGFP* expression is driven by an SV40 promoter. The self-splicing intron interrupts the reporter gene in the same direction as Alu transcription. AluY transcription is driven by its internal promoter and enhanced by an upstream 7SL enhancer. Two TTTT motifs downstream of A44 serve as RNA polymerase III transcription termination signals. The group I intron is removed by self-splicing from the Alu reporter transcript. ORF2p-mediated retrotransposition of the spliced RNA generates an intron-free *EGFP* cassette at a new genomic location, enabling *EGFP* expression. Alu, red; *EGFP*, green; self-splicing intron, gray; A44, 44-adenine Alu-derived poly(A) tract. **(b)** Base pairing schemes at the splice junctions of the wild-type group I intron and the three *EGFP*^Tet^ variants. Black arrows indicate the 5′ and 3′ splice sites. The G:U wobble base pair at the 5′ splice site and the P1, P10, and internal guide sequence (IGS) regions are shown. Lowercase letters denote exon sequences. Three versions of the *EGFP*^Tet^ reporter cassette were designed with the self-splicing intron inserted 393, 427, or 493 bp downstream of the first nucleotide of the *EGFP* start codon. Orange bases indicate the G:U wobble base pair essential for self-splicing. Magenta bases indicate nucleotide exchanges in the intron IGS or flanking *EGFP* exon sequences. **(c)** Experimental timeline and detection of self-splicing in Alu-*EGFP*^Tet^ transcripts. Cells were transfected on day 0, the culture medium was replaced, and total RNA was collected on day 4. Although Alu RNA lacks a post-transcriptionally added poly(A) tail, the encoded 3′ poly(A) tract enables oligo(dT)-primed reverse transcription. **(d)** Self-splicing of the modified group I intron. RT-PCR analysis of RNA from transfected HeLa-HA cells. The expected sizes of the cDNA products containing or lacking the self-splicing intron are ∼1.0 kb and ∼0.6 kb, respectively. *ACTB*, used as a reverse transcription control, was detected at ∼0.7 kb. Lanes “(393),” “(427),” and “(493)” correspond to pBSAluegfp(393), pBSAluegfp(427), and pBSAluegfp(493), respectively. Negative controls include water and RNA from pBSAluegfp(427)-transfected cells without reverse transcription. PCR-amplified DNA from the intron-containing pBSAluegfp(427) plasmid and the intronless pBSAluegfp(-intron) plasmid were included as controls. **(e)** Sequence analysis of self-splicing junctions. Gray boxes indicate the P1, IGS, and P10 regions. The top sequence for each construct is the reference DNA sequence. Light green regions indicate *EGFP* exons, and dark green regions indicate the self-splicing intron. The bottom sequences are cDNAs from HeLa-HA cells transfected with pBSAluegfp(393), pBSAluegfp(427), or pBSAluegfp(493), aligned with their respective reference sequences. Red dashed regions mark the sequences precisely removed by self-splicing, and the corresponding Sanger-sequencing chromatograms are shown.

Because self-splicing efficiency may depend on the surrounding exon sequence and the intron insertion site (Esnault et al., 2002), we generated three reporter cassettes, termed *EGFP*^Tet^(393), *EGFP*^Tet^(427), and *EGFP*^Tet^(493), in which the modified group I intron was inserted 393, 427, or 493 bp downstream of the first nucleotide of the *EGFP* start codon, respectively. Each cassette was positioned in the antisense orientation between the 3′ end of the AluY sequence and the encoded 44-adenine poly(A) tract, placing the group I intron in the same orientation as the Alu transcript (Figure 1a, b; pBSAluegfp[393], pBSAluegfp[427], and pBSAluegfp[493]). An upstream 7SL enhancer augmented Alu transcription by Pol III, and two TTTT motifs downstream of the encoded poly(A) tract served as Pol III transcription termination signals (Chu et al., 1995; Dewannieux et al., 2003). The reporter cassette also contained an SV40 promoter to drive EGFP expression following genomic insertion.

### Validation of self-splicing in the *EGFP*^Tet^ reporter cassette

We next tested whether the reporter constructs generated the expected intron-free products (Figure 1c). The three reporter plasmids were transfected into HeLa-HA cells, and total RNA was collected. Although Alu RNA lacks a canonical post-transcriptionally added poly(A) tail, its encoded 3′ poly(A) tract enables oligo(dT)-primed reverse transcription. The intronless plasmid pBSAluegfp(-intron) was analyzed in parallel. RT-PCR using primers flanking the intron detected the expected intron-free products for all three *EGFP*^Tet^ cassettes (Figure 1d), and Sanger sequencing confirmed precise intron removal at the predicted splice sites (Figure 1e). These results confirmed precise self-splicing in all three *EGFP*^Tet^ constructs.

### Alu-*EGFP*^Tet^ detects Alu retrotransposition in an ORF2p-dependent manner

We next asked whether the *EGFP*^Tet^ reporter could detect Alu retrotransposition in an ORF2p-dependent manner. To this end, we generated a plasmid co-expressing Alu-*EGFP*^Tet^ and monocistronic ORF2p and carrying a blasticidin resistance cassette (Figure 2a). Following transfection of Alu-permissive HeLa-HA cells (Hulme et al., 2007; Moldovan et al., 2024, 2025) and blasticidin selection, EGFP-positive cells were readily detected by flow cytometry, whereas the RT-deficient D702A mutant produced only background signals (Figure 2b). EGFP-positive cells isolated by fluorescence-activated cell sorting remained fluorescent, but the sorted EGFP-negative population and RT-deficient control population showed only background fluorescence (Figure 2c). These results demonstrated ORF2p RT-dependent fluorescence detection and fluorescence-based enrichment of reporter-positive cells.

**Figure 2.**
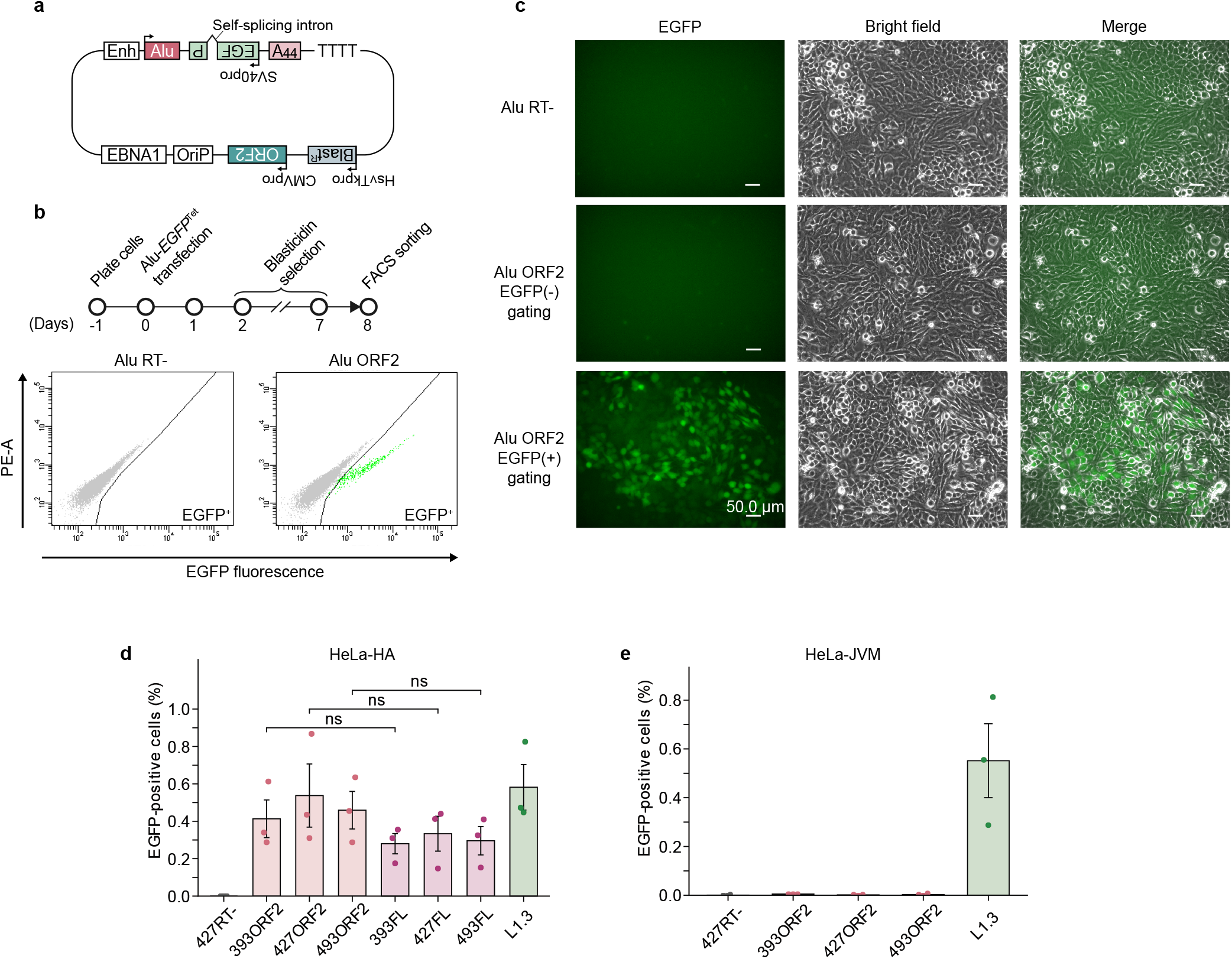
Detection and sorting of Alu-retrotransposed cells using the SINE-*EGFP*^Tet^ reporter. **(a) Schematic of the plasmid used to initiate Alu retrotransposition.** The plasmid contains the EBNA1 and oriP sequences that support episomal maintenance and a blasticidin-resistance gene for selection. It co-expresses Alu-*EGFP*^Tet^ and monocistronic ORF2p. **(b)** Fluorescence detection of Alu-retrotransposed cells by flow cytometry. (Top) Experimental timeline for the Alu-*EGFP*^Tet^ retrotransposition assay. A plasmid co-expressing the Alu-*EGFP*^Tet^(427) reporter and either wild-type ORF2p or the RT-deficient D702A mutant was transfected into HeLa-HA cells. Transfected cells were selected with blasticidin and analyzed and sorted on day 8. (Bottom) Representative FACS plots of cells expressing wild-type ORF2p (“Alu ORF2”) or the RT-deficient mutant (“Alu RT-”). The EGFP-positive gate was set using the RT-deficient control. **(c)** Fluorescence detection of Alu-retrotransposed cells by microscopy. Fluorescence microscopy of the RT-deficient control and the EGFP-negative and EGFP-positive populations sorted from cells expressing wild-type ORF2p. Sorted cells were cultured for 6 days before imaging. Fluorescence was retained in the sorted EGFP-positive population, but the EGFP-negative and RT-deficient control populations showed only background fluorescence. Scale bar, 50 µm. **(d)** Quantitative analysis of Alu retrotransposition frequencies in HeLa-HA cells. The constructs tested were Alu-*EGFP*^Tet^(393), Alu-*EGFP*^Tet^(427), and Alu-*EGFP*^Tet^(493), with retrotransposition supported by monocistronic ORF2p or full-length L1 (FL); L1.3-*mEGFPI* was included as a reporter of L1 cis retrotransposition for comparison. Alu-*EGFP*^Tet^(427) with RT-deficient ORF2p served as a negative control. **(e)** Quantitative analysis of EGFP-positive fractions in HeLa-JVM cells. The conditions tested were Alu-*EGFP*^Tet^(427) with RT-deficient ORF2p, the three Alu-*EGFP*^Tet^ reporters with wild-type ORF2p, and L1.3-*mEGFPI* for L1 cis retrotransposition. Data in (d) and (e) are mean ± SEM from n = 3 independent experiments. Corresponding monocistronic ORF2p and full-length L1 conditions in (d) were compared using two-sided Welch’s t-tests with Holm correction across the three comparisons. ns, not significant.

We then compared the three *EGFP*^Tet^ variants with monocistronic ORF2p or full-length L1 (Figure 2d). All three reporters produced detectable EGFP-positive populations. Alu-*EGFP*^Tet^(427) showed the highest average reporter-positive fraction with both the monocistronic ORF2p and full-length L1. These results are consistent with previous observations that ORF2p is sufficient to support detectable Alu mobilization (Dewannieux et al., 2003; Wallace et al., 2008).

For comparison, the established L1.3-*mEGFPI* reporter, which measures L1 cis retrotransposition using a canonical mRNA-type intron (Miyoshi et al., 2019; Ostertag et al., 2000), produced an EGFP-positive fraction of ∼0.58% in HeLa-HA cells (Figure 2d). Thus, the Alu-*EGFP*^Tet^ assay generated a readily measurable EGFP-positive fraction of the same order as that obtained with the L1 reporter.

We next tested the reporters in HeLa-JVM cells, which support L1 retrotransposition but are poorly permissive for Alu mobilization (Moldovan et al., 2024). The L1 reporter yielded a detectable EGFP-positive fraction in both cell lines; however, all three Alu-*EGFP*^Tet^ reporters produced only background-level signals in HeLa-JVM cells (Figure 2d, e). Expected intron-free products were also detected in HeLa-JVM cells (Supplementary Figure 1), suggesting that the absence of reporter-positive cells was not simply explained by failure to generate the intron-free product. Alu-*EGFP*^Tet^(427) was further used for the experiments described below. Together, these results recapitulated the previously described difference between Alu-permissive HeLa-HA and poorly permissive HeLa-JVM cells.

### Genomic insertion and target-site analysis of Alu-*EGFP*^Tet^ retrotransposition

To determine whether EGFP-positive cells harbored newly integrated Alu-*EGFP*^Tet^ sequences, EGFP-positive cells were sorted and clonally expanded. Inverse PCR and Sanger sequencing identified five independent insertions (#1-#5). Locus-specific PCR detected both an empty allele and a larger insertion-containing allele in each clone, consistent with heterozygous insertions at the respective loci (Figure 3a, b). The insertions mapped to an intron of *UBP1* on chromosome 3 (#1), an endogenous L1PA11 sequence on chromosome 12 (#2), an exon of *PPT1* on chromosome 1 (#3), an intron of *REC114* on chromosome 15 (#4), and an intron of *UST* on chromosome 6 (#5) (Figure 3c). All recovered insertions contained the full-length Alu-*EGFP*^Tet^ reporter sequence. The self-splicing intron was absent, consistent with retrotransposition of processed RNA intermediates.

**Figure 3.**
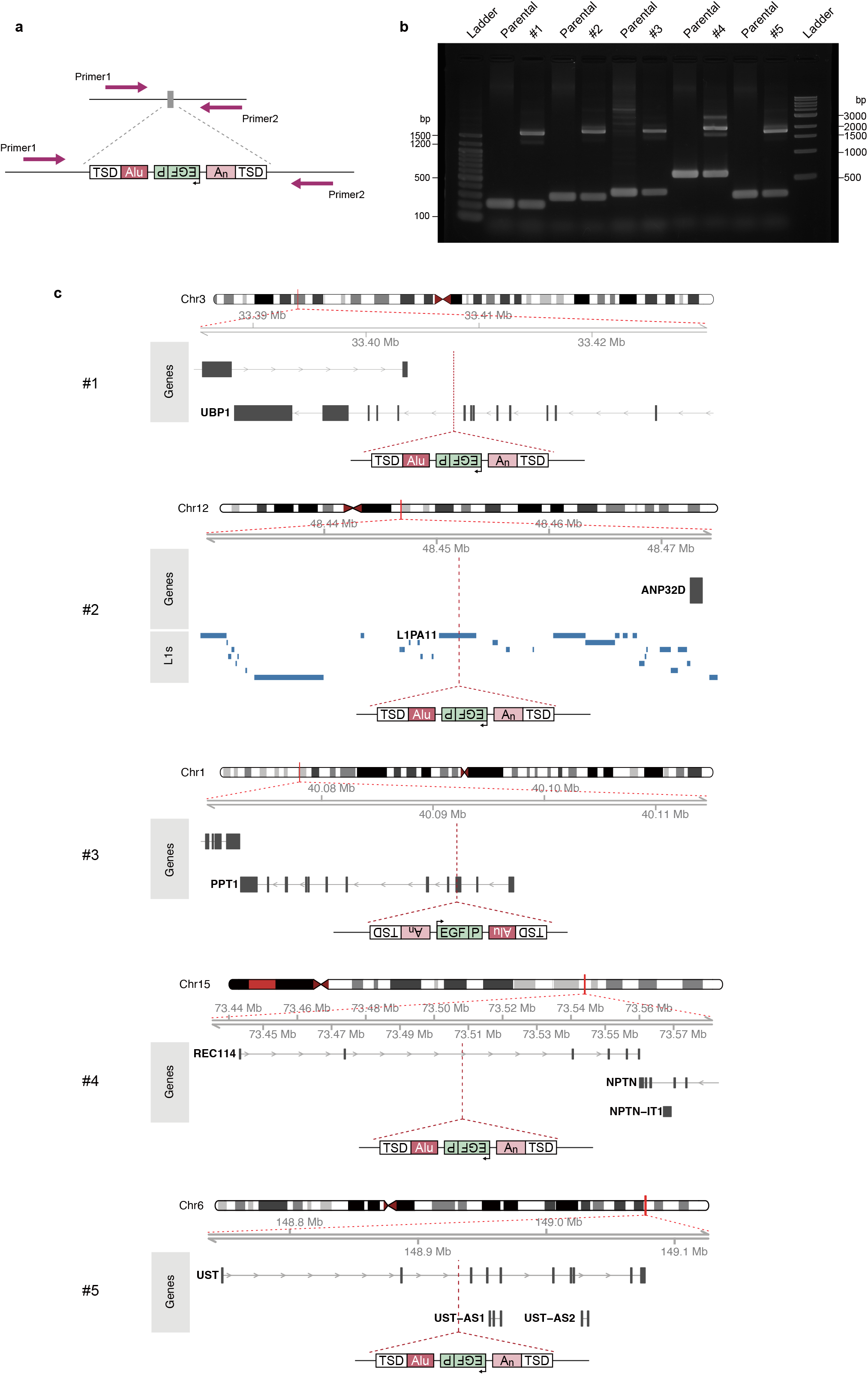
Genomic insertion and target-site analysis of Alu-*EGFP*^Tet^ via *de novo* retrotransposition. **(a) Schematic of locus-specific PCR used to detect Alu-*EGFP*^Tet^ insertion-containing alleles.** Primer pairs flanking each insertion site amplify a shorter empty-allele product and a longer product containing the Alu-*EGFP*^Tet^ insertion. **(b)** Validation of Alu insertions. Genomic DNA from control HeLa-HA cells (Parental) and from each EGFP-positive clone (#1– #5) showed a PCR product corresponding to the empty allele. Each clone also showed a longer product corresponding to the allele containing the intron-free Alu-*EGFP*^Tet^ insertion, consistent with a heterozygous insertion. **(c) *De novo* Alu insertion sites containing Alu-*EGFP*^Tet^.** The insertion sites were identified from the sequencing results using the UCSC Genome Browser BLAT tool with the GRCh38/hg38 human genome assembly. #1, #2, #3, #4, and #5 mapped to an intron of *UBP1* on chromosome 3, an endogenous L1PA11 sequence on chromosome 12, an exon of *PPT1* on chromosome 1, an intron of *REC114* on chromosome 15, and an intron of *UST* on chromosome 6, respectively. Red dashed lines indicate the genomic regions enlarged in the local view and the corresponding insertion sites, gray boxes represent exons, and blue boxes represent endogenous L1 sequences. In all five clones, the full-length Alu-*EGFP*^Tet^ reporter was inserted without the self-splicing intron and was flanked by target-site duplications (TSDs). Each insertion also retained a 3′ poly(A) tract. Detailed junction features are summarized in Table 1 and Supplementary Data 1.

**Table 1.**
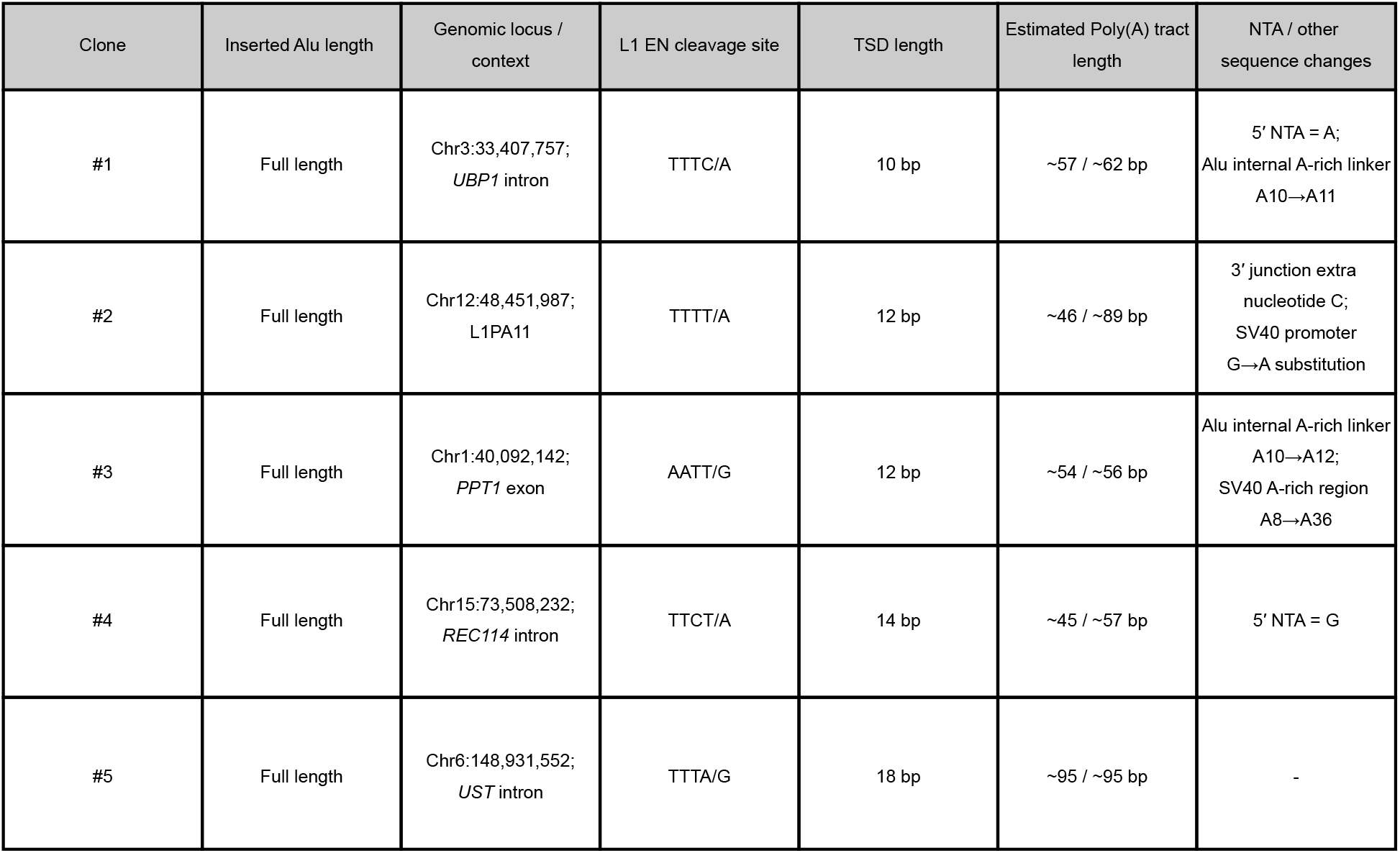
Genomic and junction features of de novo Alu*-EGFP*^Tet^ insertions. Genomic locations and insertion-junction features of five independent full-length, intron-free Alu-*EGFP*^Tet^ insertions recovered from EGFP-positive HeLa-HA clones. TSD, target-site duplication; EN, L1 endonuclease; NTA, non-templated nucleotide addition. Poly(A)-tract lengths are approximate estimates obtained by Sanger sequencing of two bacterial plasmid clones derived from a locus-specific PCR product for each insertion. Both estimates are shown; the parental reporter contained a 44-adenine poly(A) tract. A single extra nucleotide at the 3′ junction of #2 is listed separately from 5′ NTAs. The slash in each EN cleavage-site sequence indicates the inferred nick position. A dash indicates that no 5′ NTA was detected. Genomic coordinates are based on the GRCh38/hg38 human genome assembly, and gene and transcript annotations are based on NCBI RefSeq.

All five insertions showed junction features characteristic of L1-mediated TPRT (Table 1 and Supplementary Data 1). The insertions were flanked by 10–18-bp target-site duplications (TSDs), and generally occurred at AT-rich motifs preferred by the L1 EN (Feng et al., 1996; Moran et al., 1996; Wagstaff et al., 2012). Estimated 3′ poly(A)-tract lengths in all five insertions exceeded 44 nucleotides, consistent with previously reported A-tail expansion (Wagstaff et al., 2012); length increases in internal A-rich tracts were also observed in some recovered insertions (Table 1; Supplementary Data 1). Non-templated nucleotide additions (NTAs) were detected at the 5′ junctions of a subset of insertions (Baldwin et al., 2024; Gilbert et al., 2005; Moldovan et al., 2024). Detailed genomic and junction features are summarized in Table 1.

Together, the recovery of five independent, full-length, intron-free Alu-*EGFP*^Tet^ insertions with features characteristic of TPRT demonstrates that the EGFP-positive clones harbored bona fide de novo Alu retrotransposition events.

### Alu*-EGFP*^Tet^ enables quantitative analysis of positive and negative host regulators

We next tested whether *EGFP*^Tet^ could measure the effects of host factors that regulate Alu mobilization. We first examined the SRP9/14 heterodimer, a positive regulator of Alu retrotransposition (Ahl et al., 2015; Bennett et al., 2008). SRP9-targeting siRNA reduced both SRP9 and SRP14 protein levels, and SRP14-targeting siRNA similarly reduced both proteins (Figure 4a), as observed previously (Gussakovsky et al., 2023). Knockdown of MOV10, a negative regulator of Alu and L1 retrotransposition, was also confirmed by western blotting (Figure 4b).

**Figure 4.**
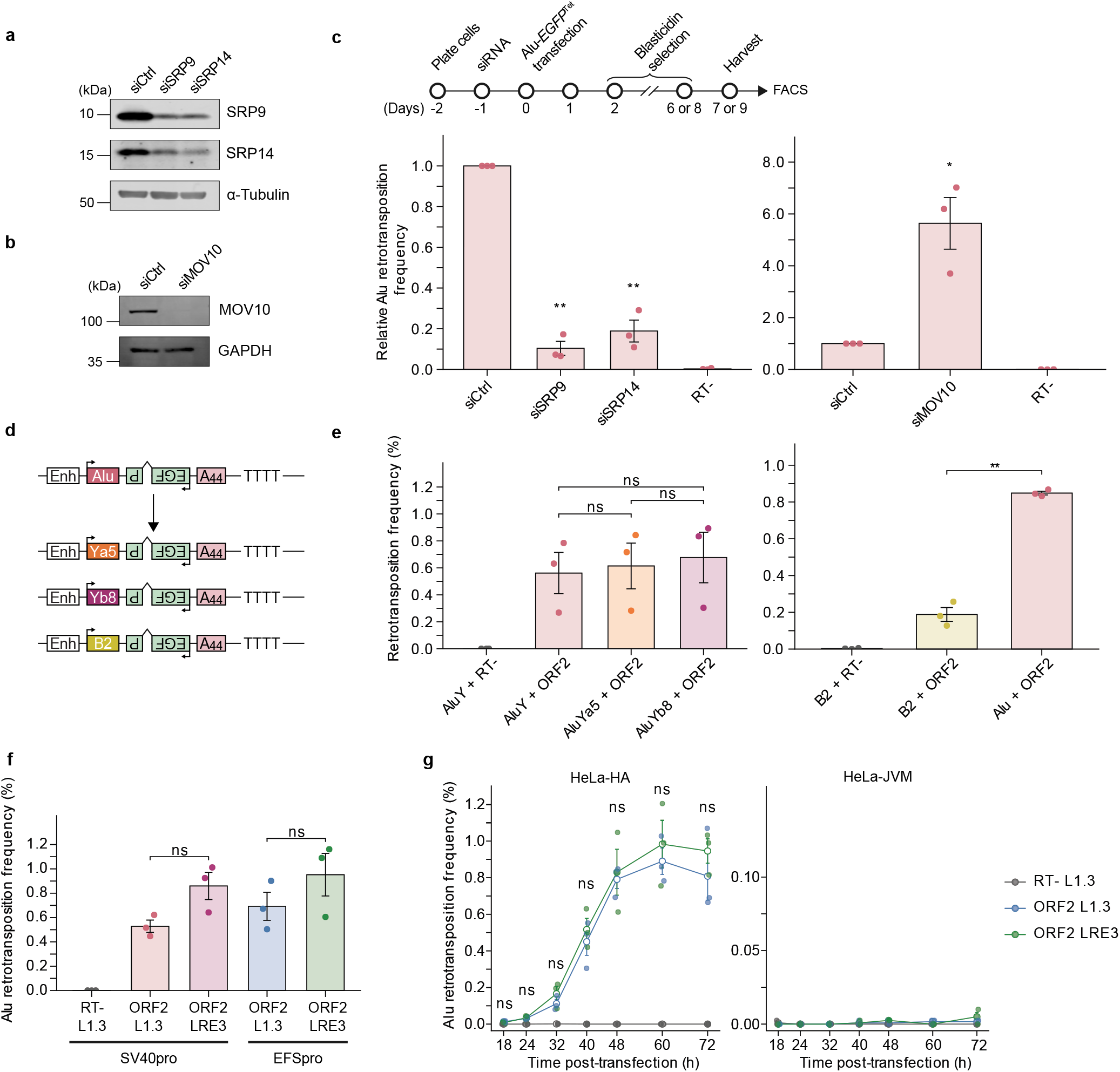
Versatility of the SINE-*EGFP*^Tet^ reporter system. **(a) siRNA knockdown of SRP9 or SRP14.** Western blot analysis of SRP9- or SRP14-knockdown cells. Introduction of siRNA targeting either SRP9 or SRP14 resulted in reduced levels of both proteins. α-Tubulin served as a loading control. **(b)** siRNA knockdown of MOV10. Western blot analysis of MOV10-knockdown cells. GAPDH served as a loading control. **(c) Quantitative analysis of Alu retrotransposition following depletion of host regulators.** (Top) Experimental timeline for Alu retrotransposition in siRNA-treated HeLa-HA cells. Cells were analyzed on day 7 for MOV10 knockdown and day 9 for SRP9/14 knockdown. (Bottom) Relative Alu retrotransposition frequency after SRP9, SRP14, or MOV10 knockdown. Values were normalized to the corresponding non-targeting siRNA control (siCtrl), which was set to 1.0. “RT-” denotes the RT-deficient ORF2p control in siCtrl-treated cells. **(d)** Schematics of AluY-, AluYa5-, AluYb8-, and B2-*EGFP*^Tet^ reporters. The AluY sequence in pBSAluegfp(427) was replaced with the indicated Alu subfamily or mouse B2 sequence, while retaining the *EGFP*^Tet^ cassette and the poly(A) tract. **(e) Detection of retrotransposition using Alu subfamily and B2-*EGFP*^Tet^ reporters.** (Left) HeLa-HA cells were co-transfected with AluY-, AluYa5-, or AluYb8-*EGFP*^Tet^ and wild-type ORF2p. The AluY reporter with RT-deficient ORF2p served as a negative control. (Right) HeLa-HA cells were co-transfected with Alu-*EGFP*^Tet^ or B2-*EGFP*^Tet^ and wild-type or RT-deficient ORF2p. **(f)** Comparison of L1.3-derived and LRE3-derived monocistronic ORF2p. Alu-*EGFP*^Tet^ reporters in which *EGFP* expression was driven by the SV40 or EFS promoter were tested with L1.3-derived or LRE3-derived ORF2p. **(g)** Time-course analysis of Alu-*EGFP*^Tet^ retrotransposition in HeLa-HA and HeLa-JVM cells. EGFP-positive fractions from Alu-EFS-*EGFP*^Tet^ were measured at 18, 24, 32, 40, 48, 60, and 72 hours after transfection with RT-deficient L1.3-derived ORF2p, wild-type L1.3-derived ORF2p, or wild-type LRE3-derived ORF2p. No drug selection was applied. Data in (c) and (e–g) are mean ± SEM from n = 3 independent experiments. Statistical analyses were two-sided one-sample t-tests in (c), with Holm adjustment for the SRP9 and SRP14 comparisons; Welch’s one-way ANOVA followed by Games–Howell tests for the Alu-subfamily comparison in (e); two-sided Welch’s t-tests for the Alu–B2 comparison in (e); two-sided Welch’s t-tests with Holm adjustment for the L1.3–LRE3 comparisons in (f); and two-sided Welch’s t-tests with Holm adjustment across the seven time points for the L1.3–LRE3 comparison in HeLa-HA cells in (g). ns, not significant; *P < 0.05; **P < 0.01.

Under SRP9 or SRP14 knockdown conditions, Alu retrotransposition measured by Alu-*EGFP^T^*^et^ decreased to ∼10–20% of the control level (Figure 4c), supporting an important role for the SRP9/14 heterodimer in efficient Alu mobilization. In contrast, MOV10 knockdown increased Alu-*EGFP*^Tet^ retrotransposition ∼5.6-fold relative to the control siRNA condition (Figure 4c), consistent with a restrictive role for MOV10 (Arjan-Odedra et al., 2012; Goodier et al., 2012). Thus, *EGFP*^Tet^ can quantify the effects of both positive and negative host regulators on Alu retrotransposition.

### *EGFP*^Tet^ is applicable across active Alu subfamilies and to a tRNA-derived SINE

To examine whether *EGFP*^Tet^ could be applied to distinct active human Alu subfamilies, we generated additional *EGFP*^Tet^ reporters carrying AluYa5 or AluYb8 sequences and compared them with the AluY reporter in the same vector backbone (Figure 4d). The three reporters yielded EGFP-positive fractions of the same order under the same ORF2p-driven assay conditions (Figure 4e), indicating that *EGFP*^Tet^ is compatible with multiple active Alu subfamilies.

To test whether *EGFP*^Tet^ could be extended beyond 7SL-derived Alu elements, we replaced Alu with the mouse B2 SINE (Figure 4d). B2 elements are also non-autonomous retrotransposons transcribed by Pol III and mobilized by ORF2p; however, they are derived from tRNA rather than 7SL RNA (Dewannieux & Heidmann, 2005a; Kroutter et al., 2009; Moldovan et al., 2024). B2-*EGFP*^Tet^ produced an EGFP-positive fraction of ∼0.2% with wild-type human ORF2p but only background signal with RT-deficient ORF2p (Figure 4e), showing that *EGFP*^Tet^ is not restricted to Alu and can detect mobilization of a phylogenetically distinct Pol III-transcribed SINE. Under the same assay conditions, B2-*EGFP*^Tet^ produced a lower reporter-positive fraction than Alu-*EGFP*^Tet^ (Figure 4e).

We also used *EGFP*^Tet^ to compare Alu trans-mobilization supported by ORF2p from two distinct human L1 elements, L1.3 and LRE3 (Farley et al., 2004). With EGFP expression driven by either the SV40 or EFS promoter, LRE3-derived ORF2p produced a higher average reporter-positive fraction than L1.3-derived ORF2p, although neither comparison was statistically significant (Figure 4f).

### Time-course analysis of Alu-*EGFP*^Tet^ retrotransposition

We next performed a time-course analysis to determine when Alu-*EGFP*^Tet^-positive cells became detectable (Figure 4g). In HeLa-HA cells, EGFP-positive cells were detected by ∼24 hours after transfection, increased between 32 and 48 hours, and approached a plateau between 48 and 60 hours. Early Alu retrotransposition was also detected in previous colony-based assays (Kroutter et al., 2009). L1.3- and LRE3-derived ORF2p showed similar time courses, while the RT-deficient control showed no appreciable increase above background.

In HeLa-JVM cells, Alu reporter activity remained near background at every time point with wild-type ORF2p expression. Thus, the difference between HeLa-HA and HeLa-JVM cells persisted throughout the assay window and did not support a model in which EGFP-positive cells arose early in HeLa-JVM cells and were subsequently lost. Low Alu reporter activity in HeLa-JVM cells was observed with both SV40- and EFS-driven *EGFP* expression and with ORF2p derived from either L1.3 or LRE3 (Figures 2e and 4g).

### Fluorescent variants extend *EGFP*^Tet^ for multicolor detection

Because self-splicing depends on interactions between the intron and flanking exon sequences, generating another SINE reporter with a different coding sequence may require redesign of the exon-intron junction. To enable multicolor analysis, we instead converted *EGFP*^Tet^ stepwise to the cyan variants *ECFP*^Tet^, *Cerulean*^Tet^, and *mTurquoise2*^Tet^ through defined amino acid substitutions (Figure 5a) (Goedhart et al., 2012; Heim & Tsien, 1996; Rizzo et al., 2004; Tsien, 1998). We also replaced the SV40 promoter with the EFS promoter to generate Alu-EFS-*mTurquoise2*^Tet^. The mTurquoise2 derivatives produced the highest average cyan-positive fractions among the tested variants (Figure 5b). Only background signal was detected with Alu-EFS-*mTurquoise2*^Tet^ and RT-deficient ORF2p.

**Figure 5.**
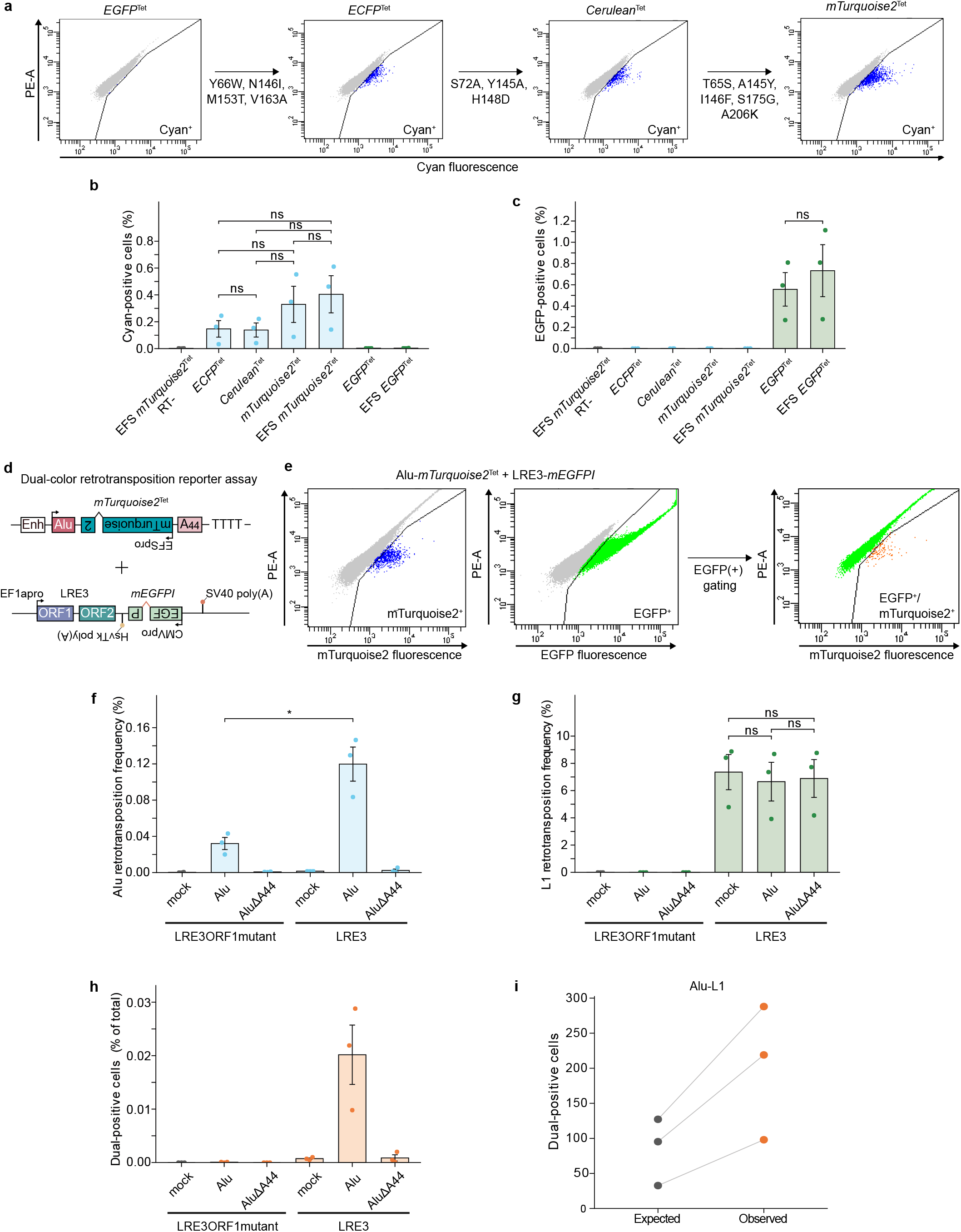
Fluorescent variants of *EGFP*^Tet^ and dual-color detection of Alu trans-mobilization and L1 cis retrotransposition. **(a) Sequential conversion of *EGFP*^Tet^ to *ECFP*^Tet^, *Cerulean*^Tet^, and *mTurquoise2*^Tet^.** An *ECFP*^Tet^ reporter was generated by introducing four amino acid substitutions (Y66W, N146I, M153T, and V163A) into the *EGFP* sequence of the original Alu-*EGFP*^Tet^ reporter plasmid. Additional amino acid substitutions (S72A, Y145A, and H148D) in *ECFP*^Tet^ generated *Cerulean*^Tet^, which was further converted to an *mTurquoise2*^Tet^ reporter by introducing T65S, A145Y, I146F, S175G, and A206K. Representative flow-cytometric plots are shown. **(b)** Cyan-positive fractions measured using the cyan detection channel. The y-axis represents the percentage of cyan-positive cells. The conditions tested in HeLa-HA cells were EFS-*mTurquoise2*^Tet^ with RT-deficient ORF2p and *ECFP*^Tet^, *Cerulean*^Tet^, *mTurquoise2*^Tet^, EFS-*mTurquoise2*^Tet^, *EGFP*^Tet^, or EFS-*EGFP*^Tet^ with wild-type ORF2p. **(c)** EGFP-positive fractions measured in the same reporter series. All conditions in (b), including *EGFP*^Tet^ and EFS-*EGFP*^Tet,^ were measured in the EGFP detection channel. **(d)** Schematic of the dual-color retrotransposition reporter assay. Alu-EFS-*mTurquoise2*^Tet^ was combined with an *mEGFPI*-tagged LRE3 reporter to detect Alu trans-mobilization and L1 cis retrotransposition within the same cell population. The reporter conditions consisted of pBSmock, intact Alu-EFS-*mTurquoise2*^Tet^, or Alu-Δpoly(A)-*mTurquoise2*^Tet^ combined with wild-type or ORF1p R261A/R262A-mutant LRE3-*mEGFPI*. ΔA44 denotes deletion of the encoded 44-adenine poly(A) tract. **(e)** Representative gating of mTurquoise2-positive, EGFP-positive, and dual-positive cells. Before fluorescence gating, cells were gated using SSC-A versus FSC-A, and singlets were selected using sequential FSC-W vs FSC-H and SSC-W vs SSC-H gating. mTurquoise2-positive and EGFP-positive gates were then defined using the corresponding negative controls and conditions in which only one reporter was detectably active, and EGFP/mTurquoise2 dual-positive events were identified. Representative plots from HeLa-HA cells are shown. The rightmost plot shows mTurquoise2 fluorescence among EGFP-positive events. The complete gating strategy and representative control plots are shown in Supplementary Figure 2. **(f)** mTurquoise2-positive fractions under wild-type or ORF1p-mutant L1 expression. Mock, intact Alu-EFS-*mTurquoise2*^Tet^, and Alu-Δpoly(A)-*mTurquoise2*^Tet^-transfected HeLa-HA cells were examined with wild-type or ORF1p-mutant LRE3-*mEGFPI*. **(g)** L1-derived EGFP-positive fractions under the same conditions. L1 retrotransposition with mock, intact Alu, and Alu-Δpoly(A) constructs under wild-type or ORF1p-mutant L1 backgrounds was detected using the EGFP detection channel. The same samples as in (f) were analyzed. **(h)** Analysis of EGFP/mTurquoise2 dual-positive cells under the same conditions. Dual-positive events were defined as events within the EGFP-positive gate that exceeded the mTurquoise2 threshold. Dual-positive fractions were calculated relative to the total number of analyzed singlet events. The same samples as in (f) were analyzed. **(i)** Observed and expected dual-positive event counts in the Alu–L1 assay. For each biological replicate with wild-type L1 and intact Alu-EFS-*mTurquoise2*^Tet^, event counts from two technical replicates containing 500,000 singlet events each were pooled and are shown per 1,000,000 singlet events. Expected dual-positive counts under independence were calculated as (mTurquoise2-positive events × EGFP-positive events) / total singlet events. Lines connect observed and expected counts from the same biological replicate. The three independent experiments are shown individually. Observed-to-expected ratios are provided in Supplementary Table 1. Data in (b), (c) and (f–h) are mean ± SEM from n = 3 independent experiments. Statistical analyses were Welch’s one-way ANOVA followed by Games–Howell tests for the four cyan reporter variants in (b) and the mock, intact Alu, and poly(A)-deficient Alu groups with wild-type L1 in (g), and two-sided Welch’s t-tests for *EGFP*^Tet^ versus EFS-*EGFP*^Tet^ in (c) and intact Alu with wild-type versus ORF1p-mutant L1 in (f). ns, not significant; *P < 0.05.

The cyan and EGFP-based reporters showed minimal signal in the EGFP and cyan detection channels, respectively (Figure 5b, c), demonstrating reciprocal channel separation under these assay conditions. Because the reporters differ in fluorophore properties and, for the EFS construct, promoter activity, the measured reporter-positive fractions should not be interpreted as direct differences in the underlying Alu retrotransposition frequency. Thus, the EGFP reporter could be converted to cyan without redesigning the self-splicing exon-intron junctions. Based on its cyan-positive fraction and channel separation from EGFP, Alu-EFS-*mTurquoise2*^Tet^ was selected for subsequent dual-color experiments.

### Dual-color analysis enables concurrent detection of Alu trans-mobilization and L1 cis retrotransposition at single-cell resolution

To our knowledge, Alu trans-mobilization and L1 cis retrotransposition have not previously been monitored concurrently within the same cell population at single-cell resolution. Because Alu and L1 share ORF2p, we examined both the effect of Alu co-expression on L1 retrotransposition and the distribution of the two reporter signals across individual cells. We therefore combined Alu-EFS-*mTurquoise2*^Tet^ with a full-length L1 retrotransposition reporter based on the highly active human LRE3 element and carrying the *mEGFPI* cassette (LRE3-*mEGFPI*) to monitor Alu trans-mobilization and L1 cis retrotransposition in the same cell population (Figure 5d, e; Supplementary Figure 2a-e). Wild-type LRE3-*mEGFPI* was compared with an otherwise identical ORF1p-mutant reporter carrying the R261A/R262A substitutions (Figure 5f-h) (Martin et al., 2005; Moran et al., 1996; Wei et al., 2001). The mutant retained the ORF2 coding sequence but produced only a background-level L1 reporter-positive fraction. This design enabled examination of Alu trans-mobilization when L1 cis retrotransposition was largely abolished. Each L1 reporter was co-transfected with intact Alu-EFS-*mTurquoise2*^Tet^, Alu-Δpoly(A)-*mTurquoise2*^Tet^, or mock.

The intact Alu reporter yielded cyan-positive fractions of ∼0.032% with the ORF1p-mutant L1 reporter and ∼0.12% with the wild-type L1 reporter (Figure 5f). Deletion of the Alu poly(A) tract reduced the cyan-positive fraction to near-background levels with either L1 reporter. Thus, wild-type L1 supported an approximately fourfold higher Alu retrotransposition frequency than its ORF1p-mutant counterpart, while low-frequency, poly(A)-dependent Alu retrotransposition remained detectable with the ORF1p-mutant L1.

In contrast, the EGFP-positive fraction from the wild-type L1 reporter was ∼7% after co-transfection with mock, intact Alu, or poly(A)-deficient Alu, with no statistically significant difference among the three groups (Figure 5g). The ORF1p-mutant L1 reporter produced only background-level EGFP signal, consistent with a previous report (Moran et al., 1996). Thus, neither Alu reporter co-transfection nor poly(A)-dependent Alu mobilization significantly reduced L1 retrotransposition under these conditions.

With the wild-type L1 reporter and intact Alu-EFS-*mTurquoise2*^Tet^, an EGFP/mTurquoise2 dual-positive population was reproducibly detected in all three independent experiments and averaged ∼0.020% of singlet events (Figure 5e, h). Dual-positive fractions in the mock, poly(A)-deficient Alu, and ORF1p-mutant L1 controls were near background (Supplementary Table 2 and Supplementary Figure 2c-e).

We next compared the observed dual-positive counts with those expected if Alu-positive and L1-positive cells were independently distributed. The observed dual-positive counts exceeded the expected counts in all three experiments, with observed-to-expected ratios of 2.3–3.0 (Figure 5i and Supplementary Table 1).

As a complementary test, we co-transfected Alu-*EGFP*^Tet^ and Alu-EFS-*mTurquoise2*^Tet^ in the presence of monocistronic ORF2p (Supplementary Figure 3a–d). The average Alu-*EGFP*^Tet^-positive fraction was lower with intact or poly(A)-deficient cyan Alu than with mock, but the differences were not statistically significant. Observed dual-positive counts exceeded independence-based expectations in all three experiments with two intact Alu reporters (observed-to-expected ratios, 6.8–33.4; Supplementary Figure 3e and Supplementary Table 1). Taken together, dual-positive cells occurred more frequently than expected under independence in both the Alu–L1 and Alu–Alu assays.

## Discussion

We developed *EGFP*^Tet^ and its cyan fluorescent derivative to detect retrotransposition of Pol III-transcribed SINEs by fluorescence. *EGFP*^Tet^ builds on the group I self-splicing intron system established in *neo*^Tet^, extending this approach to fluorescence-based assays for flow-cytometric quantification, microscopy, cell sorting, time-course analysis, and multicolor analysis. Precise self-splicing and recovery of full-length, intron-free Alu-*EGFP*^Tet^ insertions bearing TSDs, L1 EN-preferred cleavage sites, and 3′ poly(A) tracts demonstrate bona fide de novo Alu retrotransposition in reporter-positive clones.

Time-course analysis of Alu-*EGFP*^Tet^ also reproduced the known difference in Alu permissiveness between HeLa-HA and HeLa-JVM cells (Moldovan et al., 2024), arguing against transient early activation of the reporter followed by selective loss of reporter-positive cells in HeLa-JVM cells. Instead, differences in RNA metabolism, host-factor activity, or other intracellular features may contribute to this cell-line-specific Alu permissiveness. This assay also revealed host-factor-dependent regulation: SRP9/14 knockdown reduced Alu retrotransposition frequency, but MOV10 knockdown increased it. Thus, *EGFP*^Tet^ provides a fluorescence-based assay for analyzing host regulators of Alu mobilization and may be suitable for genetic or chemical screening.

The Alu subfamily, B2, and ORF2p-comparison experiments further show that *EGFP*^Tet^ can be used to compare different SINE substrates and ORF2p sources. Future assays pairing mouse B2 with mouse L1 ORF2p could test species-specific compatibility between SINE substrates and LINE proteins, and extension to animal models may enable analysis of tissue- or developmental-stage-specific SINE mobilization. Application to the independently evolved, Alu-like dimeric 7SL-derived SINE Urop from the shrew mole (Kosushkin et al., 2026) could help define RNA features that permit access to the L1 machinery.

The cyan derivative was generated without redesigning the self-splicing junctions, enabling Alu trans-mobilization and L1 cis retrotransposition to be distinguished within the same cell population. This makes it possible to examine not only the frequency of each type of retrotransposition, but also whether the corresponding reporter signals occur in the same cells. Wild-type L1 supported more Alu retrotransposition than the ORF1p-mutant construct, consistent with reports that ORF1p can enhance Alu mobilization, although it is not essential (Dewannieux et al., 2003; Wallace et al., 2008). However, ORF2p abundance and RNP assembly were not compared between the two constructs, so the difference cannot be attributed specifically to a direct effect of ORF1p on Alu mobilization. The ORF1p-mutant L1 supported Alu mobilization despite its own retrotransposition defect, suggesting that endogenous L1 copies that cannot retrotranspose themselves could still support Alu mobilization if they express functional ORF2p (Belancio et al., 2010).

Although previous work provided evidence for competition between L1 and Alu RNAs for ORF2p (Doucet et al., 2015), Alu co-expression did not significantly reduce L1 retrotransposition in our assay. Notably, Alu and L1 reporter signals co-occurred more frequently than expected under independence in all three experiments. These observations do not exclude molecular competition, but they do not support a simple model in which Alu mobilization is accompanied by a measurable reduction in L1 retrotransposition under these conditions. An excess of dual-positive cells was also observed in the complementary Alu–Alu assay. Shared cellular permissiveness, including differences in cell-cycle state, gene expression, or host factor availability, could contribute to this pattern, although sequential use of a shared ORF2p-containing machinery or facilitation of one event by another remains possible. Law and Burns identified chimeric L1 insertions involving diverse RNAs, including Alu sequences (Law & Burns, 2026). Concurrent L1 insertions have also been implicated in reciprocal translocations and complex chromosomal rearrangements in human tumors, underscoring the importance of investigating multiple retrotransposition events and their genomic consequences within individual cells (Zumalave et al., 2026). Clonal isolation of dual-positive cells and characterization of Alu and L1 insertion junctions would clarify the genomic structures underlying reporter co-occurrence within individual cell lineages.

This reporter has several limitations. A dedicated polyadenylation signal was omitted from the fluorescent-protein reporter to avoid potential premature Pol III termination; reporter expression after integration may therefore depend on surrounding genomic sequences. Incomplete self-splicing may also lead to under-detection; group I intron variants with higher splicing-dependent reporter activity could improve sensitivity (Hasegawa et al., 2003). Transient transfection does not reproduce endogenous SINE expression, and cell-to-cell variation in plasmid uptake or expression may contribute to reporter co-occurrence. Endpoint fluorescence alone does not resolve the temporal order of individual insertion events or establish direct cooperation between them.

Future applications could combine *EGFP*^Tet^ or *mTurquoise2*^Tet^ with temporal sorting, live-cell imaging, stable single-copy reporters, organoids, or animal models to link SINE mobilization to cell state, developmental history, and tissue environment. Evidence consistent with Alu mobilization during gametogenesis (Feusier et al., 2019), together with a proposed model linking heritable L1 insertions to early embryogenesis (de los Rios Barreda et al., 2026), suggests that the two elements may favor different developmental windows despite sharing ORF2p. Dual-color analysis in appropriate developmental models could clarify when cells support one or both processes. By making Pol III-transcribed SINE mobilization directly visible, sortable, and multiplexable, *EGFP*^Tet^ provides a route to move beyond population-averaged assays and ask when, where, and in which cellular states SINEs gain access to the L1 machinery.

## Materials and Methods

### Cell lines and cell culture conditions

HeLa-HA cells were provided by Astrid M. Roy-Engel at Tulane University, and HeLa-JVM cells were provided by John V. Moran at the University of Michigan. HeLa-HA cells were cultured in Minimum Essential Medium (MEM; Gibco or Nacalai Tesque) supplemented with 1× MEM Non-Essential Amino Acids Solution (NEAA; Nacalai Tesque), 10% (v/v) fetal bovine serum (FBS; Capricorn Scientific or MP Biomedicals), 2 mM L-glutamine (Sigma-Aldrich), 100 U/mL penicillin, and 100 µg/mL streptomycin (Sigma-Aldrich). HeLa-JVM cells were maintained in Dulbecco’s Modified Eagle’s Medium (DMEM; Shimadzu Diagnostics) supplemented with 0.165% (w/v) NaHCO₃, 10% (v/v) FBS, 2 mM L-glutamine, 100 U/mL penicillin, and 100 µg/mL streptomycin. Both cell lines were cultured at 37°C in a humidified incubator with 5% CO₂.

All cell lines were routinely tested for *Mycoplasma* contamination using the VenorGeM Classic Mycoplasma Detection Kit (Sigma-Aldrich) and confirmed to be negative. The identity of the HeLa-HA and HeLa-JVM cell lines was verified by short tandem repeat (STR) profiling.

### Detection of self-splicing of the Alu-*EGFP*^Tet^ reporters

HeLa-HA or HeLa-JVM cells were seeded in 6-well plates at 4.0 × 10^5^ cells per well. Twenty-four hours after seeding (day 0), cells were transfected with one of the following plasmids: pBSAluegfp(393), pBSAluegfp(427), pBSAluegfp(493), or the intronless control plasmid, pBSAluegfp(-intron). For each transfection, a mixture was prepared by incubating 1.0 µg of plasmid DNA with 3.0 µL of FuGENE HD (Promega) in 100 µL of Opti-MEM (Thermo Fisher Scientific) for 10 minutes at room temperature. The mixture was then added dropwise to the cells.

Beginning on day 1, the culture medium was replaced daily for four days. On day 4, cells were harvested using 0.25% Trypsin-EDTA (Gibco). The detached cells were suspended in fresh MEM and transferred to 1.5 mL microcentrifuge tubes. Cells were then pelleted by centrifugation, washed with ice-cold 1× PBS (Shimadzu Diagnostics), and flash-frozen in liquid nitrogen. Cell pellets were stored at -80°C until RNA extraction. RNA was extracted and purified using the RNeasy Plus Mini Kit (Qiagen) with on-column DNase treatment (Qiagen), following the manufacturer’s protocol. Total RNA was eluted in 50 µL of RNase-free water, and its concentration was measured with a NanoDrop (Thermo Fisher Scientific).

Total RNA (1 µg) was reverse-transcribed into cDNA in a 20 µL reaction containing 0.2 mM dNTPs (TOYOBO), 1 U/µL Ribonuclease Inhibitor (TaKaRa Bio), 0.25 U/µL AMV Reverse Transcriptase XL (TaKaRa Bio), and 0.125 µM oligo(dT) primer (Invitrogen), following the manufacturer’s protocol. A no-template control (NTC), in which the RNA template was replaced with RNase-free water, and a no-reverse-transcriptase control were included. Thermal cycling conditions were 30°C for 10 minutes, 42°C for 30 minutes, and 99°C for 5 minutes. After the reaction, the cDNA was diluted with 80 µL of RNase-free water and stored at -80°C until use.

To confirm that the self-splicing intron had been spliced, PCR was performed using 5 µL of the cDNA solution as a template in a 15 µL reaction containing 7.5 µL of 2× GoTaq Green Master Mix (Promega), 0.267 µM of each primer, and nuclease-free water. The PCR program consisted of an initial denaturation at 95°C for 2 minutes; 25 cycles of denaturation at 95°C for 30 seconds, primer annealing at 55°C for 30 seconds, and extension at 72°C for 1 minute; and a final extension at 72°C for 5 minutes. The *EGFP* sequence was amplified with primers KO87 and KO88, while *ACTB* (β-actin) cDNA was amplified with primers KO69 and KO70 as an internal control. pBSAluegfp(427) and pBSAluegfp(-intron) plasmids served as positive controls for the unspliced and spliced products, respectively. PCR products were separated by electrophoresis on a 1.5% (w/v) agarose gel (Nippon Gene), stained with GelRed (Biotium), and visualized under UV light using an E-box CX5 imager (Vilber Lourmat).

To analyze the sequence of the self-spliced junctions, the *EGFP* fragments corresponding to the spliced product were excised from the gel and transferred to a 1.5 mL tube. The gel slices were dissolved in 450 µL of Buffer QG (QIAGEN) at 55°C for 10 minutes. After adding 150 µL of 2-propanol, the solution was applied to an Econospin IIα column (Gene Design) for purification. After washing the column with 500 µL of Buffer PE (QIAGEN), the column was transferred to a new 1.5 mL tube. The purified DNA was eluted with 30 µL of Elution Solution (Sigma-Aldrich) by centrifugation at ∼9,000 × g for 3 minutes. The resulting *EGFP* fragments were then subjected to Sanger sequencing using the KO87 primer.

### Alu, B2, and L1 retrotransposition assays

For retrotransposition assays, HeLa-HA or HeLa-JVM cells were seeded in 6-well plates at 1.5 × 10^5^ cells per well. Approximately 24 hours later (day 0), cells were transfected with one of the following plasmids: cepB-Alu-gfp(427)-O2F3RT(-), cepB-Alu-gfp(393)-O2F3, cepB-Alu-gfp(427)-O2F3, cepB-Alu-gfp(493)-O2F3, cepB-Alu-gfp(393)-L1F3, cepB-Alu-gfp(427)-L1F3, cepB-Alu-gfp(493)-L1F3, or cepB-gfp-L1.3. For each well, a transfection mixture containing 1.0 µg of plasmid DNA and 3.0 µL of FuGENE HD (Promega) in 100 µL of Opti-MEM was prepared, incubated for 10 minutes at room temperature, and then added to the cells. Each transfection was performed in technical duplicate. Twenty-four hours post-transfection (day 1), the medium was replaced. On day 2, selection was initiated by culturing the cells in medium containing 10 µg/mL Blasticidin S (Kaken Seiyaku). On day 8, surviving cells were harvested by trypsinization, washed with 1× PBS, and analyzed by flow cytometry. The percentage of EGFP-positive cells was determined from a total of 20,000 events per technical replicate using a BD FACSCalibur.

To examine Alu retrotransposition efficiency under SRP9/14 or MOV10 knockdown conditions, HeLa-HA cells were seeded in 6-well plates with 2 mL of medium at 1.5 × 10^5^ cells per well on day -2, as described above. On day -1, 1 mL of medium was removed from each well, and cells were transfected with siRNAs. Briefly, 1 µL of 20 µM siRNA solution (targeting SRP9 [Horizon/Dharmacon, L-019731-01-0005], SRP14 [Horizon/Dharmacon, L-017767-01-0005], MOV10 [Horizon/Dharmacon, L-014162-00-0005], or a non-targeting control [Horizon/Dharmacon, D-001810-10-0020]) was mixed with 3 µL of Lipofectamine RNAiMAX (Invitrogen) in 100 µL of Opti-MEM, incubated for 5 minutes at room temperature, and then added to the cells. Twenty-four hours after siRNA transfection (day 0), cells were transfected with 1 µg of cepB-Alu-gfp(427)-O2F3RT(-) or cepB-Alu-gfp(427)-O2F3. Plasmid transfection was performed using FuGENE HD as described previously. On day 1, the medium was replaced. Selection with 10 µg/mL Blasticidin S was initiated on day 2, and cells were analyzed on day 7 (MOV10 knockdown) or day 9 (SRP9/14 knockdown). The percentage of EGFP-positive cells was determined from 200,000 events per technical replicate using a BD FACSCanto II.

For the B2 retrotransposition assay, HeLa-HA cells were seeded in 6-well plates as described above. Twenty-four hours later (day 0), 0.5 µg of pBSB2egfp(427) and 0.5 µg of either pTMO2F3 or pTMO2F3D702A were combined with 3.0 µL of FuGENE HD and 100 µL of Opti-MEM. Transfection was performed using FuGENE HD as described previously. As a control, a mixture of 0.5 µg of pBSAluegfp(427) and 0.5 µg of pTMO2F3 was transfected using the same procedure. On day 1, the medium was replaced with fresh MEM. From day 2 to day 8, transfected cells were selected with MEM containing 200 µg/mL Hygromycin B (Fujifilm Wako). On day 9, cells were analyzed. The percentage of EGFP-positive cells was determined from 20,000 events per technical replicate using a BD FACSCanto II.

For the AluYa5 and AluYb8 retrotransposition assay, HeLa-HA cells were seeded in 6-well plates as described above. Twenty-four hours later (day 0), 0.5 µg of either pBSAluYa5egfp(427) or pBSAluYb8egfp(427) was combined with 0.5 µg of pEF1O2F3_B and mixed with 3.0 µL of FuGENE HD in 100 µL of Opti-MEM. Transfection was performed using FuGENE HD as described previously. As a control, a mixture of 0.5 µg of pBSAluegfp(427) and 0.5 µg of either pEF1O2F3_B or pEF1O2F3D702A_B was transfected using the same procedure. On day 1, the medium was replaced with fresh MEM culture medium. From day 2 to day 7, transfected cells were selected with MEM containing 10 µg/mL Blasticidin S. On day 8, cells were analyzed. The percentage of EGFP-positive cells was determined from 200,000 events per technical replicate using a BD FACSCanto II.

For the retrotransposition assay comparing L1.3-derived and LRE3-derived ORF2p, HeLa-HA cells were seeded in 6-well plates as described above. Twenty-four hours later (day 0), 0.5 µg of either pBSAluegfp(427) or pBSAluEFSegfp(427) was combined with 0.5 µg of either pEF1O2F3_B or pEF1O2F3LRE3_B, and the mixture was combined with 3.0 µL of FuGENE HD and 100 µL of Opti-MEM. Plasmid transfection was performed using FuGENE HD as described previously. As a control, a mixture of 0.5 µg of pBSAluegfp(427) and 0.5 µg of pEF1O2F3D702A_B was transfected using the same procedure. On day 1, the medium was replaced with fresh MEM. From day 2 to day 7, transfected cells were selected with MEM containing 10 µg/mL Blasticidin S. On day 8, cells were analyzed. The percentage of EGFP-positive cells was determined from 200,000 events per technical replicate using a BD FACSCanto II.

For the time-course analysis of Alu retrotransposition, HeLa-HA or HeLa-JVM cells were seeded in 6-well plates as described above. Twenty-four hours later (day 0), 0.5 µg of pBSAluEFSegfp(427) was combined with 0.5 µg of either pEF1O2F3_B or pEF1O2F3LRE3_B, and the mixture was combined with 3.0 µL of FuGENE HD and 100 µL of Opti-MEM. Plasmid transfection was performed using FuGENE HD as described previously. As a control, a mixture of 0.5 µg of pBSAluEFSegfp(427) and 0.5 µg of pEF1O2F3D702A_B was transfected using the same procedure. At 18 hours after transfection, the medium was replaced with the respective fresh culture medium. No drug selection was applied. For each condition, separate wells were used for each time point; no individual well was analyzed repeatedly over time. Cells were analyzed at 18, 24, 32, 40, 48, 60, and 72 hours after transfection. The percentage of EGFP-positive cells was determined from 20,000 events per technical replicate using a BD FACSCanto II.

For the retrotransposition assay using *ECFP*^Tet^, *Cerulean*^Tet^, and *mTurquoise2*^Tet^, HeLa-HA cells were seeded in 6-well plates as described above. Twenty-four hours later (day 0), 0.5 µg of one of the following reporter plasmids—pBSAluecfp(427), pBSAlucerulean(427), pBSAlumTurquoise2(427), pBSAluEFSmTurquoise2(427), pBSAluegfp(427), or pBSAluEFSegfp(427)—was combined with 0.5 µg of pEF1O2F3_B, then mixed with 3.0 µL of FuGENE HD and 100 µL of Opti-MEM. Transfection was performed with FuGENE HD as described previously. As a control, a mixture of 0.5 µg of pBSAluEFSmTurquoise2(427) and 0.5 µg of pEF1O2F3D702A_B was transfected using the same procedure. On day 1, the medium was replaced with fresh MEM. From day 2 to day 7, transfected cells were selected with MEM containing 10 µg/mL Blasticidin S. On day 8, cells were analyzed. The percentages of cyan-positive and EGFP-positive cells were determined from 200,000 events per technical replicate using a BD FACSCanto II.

For the dual-color retrotransposition assay of Alu and L1, HeLa-HA cells were seeded in 6-well plates as described above. Twenty-four hours later (day 0), 1.0 µg of one of the following plasmids—pBlueScript SK(-), pBSAluEFSmTurquoise2(427), or pBSAluΔA44EFSmTurquoise2(427)—was combined with 0.5 µg of either cep99-gfp-EF1LRE3 or cep99-gfp-EF1LRE3RR261_262AA and mixed with 4.5 µL of FuGENE HD in 100 µL of Opti-MEM. Transfection was performed using FuGENE HD as described previously. On day 1, the medium was replaced with fresh MEM. From day 2 to day 7, transfected cells were selected with MEM containing 1 µg/mL puromycin (Sigma-Aldrich). On day 8, cells were analyzed. Up to 500,000 singlet events were acquired per technical replicate using a BD FACSCanto II. The percentages of mTurquoise2-positive, EGFP-positive, and EGFP/mTurquoise2 dual-positive cells were calculated separately for each technical replicate.

For the dual-color retrotransposition assay of Alu and Alu, HeLa-HA cells were seeded in 6-well plates as described above. Twenty-four hours later (day 0), 1.0 µg of one of the following plasmids—pBlueScript SK(-), pBSAluEFSmTurquoise2(427), or pBSAluΔA44EFSmTurquoise2(427)—was combined with 0.5 µg of either cepB-Alu-gfp(427)-O2F3 or cepB-Alu-gfp(427)-O2F3RT(-) and mixed with 4.5 µL of FuGENE HD in 100 µL of Opti-MEM. Transfection was performed using FuGENE HD as described previously. On day 1, the medium was replaced with fresh MEM. From day 2 to day 7, transfected cells were selected with MEM containing 10 µg/mL Blasticidin S. On day 8, cells were analyzed. A total of 500,000 singlet events were acquired per technical replicate using a BD FACSCanto II. The percentages of mTurquoise2-positive, EGFP-positive, and EGFP/mTurquoise2 dual-positive cells were calculated separately for each technical replicate.

Flow-cytometric data were acquired and analyzed using BD FACSDiva software (version 6.1.3) for the BD FACSCanto II, BD FACSAria IIu and BD FACSAria III, or BD CellQuest Pro software (version 5.2) for the BD FACSCalibur. For dual-color analyses, mTurquoise2 and EGFP were excited using the 405-nm and 488-nm lasers and detected through 510/50-nm and 530/30-nm band-pass filters, respectively. Fluorescence collected through the 585/42-nm band-pass filter after 488-nm excitation, displayed as PE-A in FACSDiva, was recorded as the second axis used to define the fluorescence-positive gates. No fluorescence compensation matrix was applied. Reciprocal channel separation was assessed using reporter assays in which only one fluorescent reporter was detectably active (Figure 5b, c), and matched controls in the dual-color experiments showed low dual-gated event counts (Supplementary Table 2). For these analyses, cells were first gated using SSC-A versus FSC-A, followed by sequential singlet gating using FSC-W versus FSC-H and SSC-W versus SSC-H (Supplementary Figure 2). Fluorescence gates were defined using the corresponding negative controls and control conditions in which only one reporter was detectably active.

### *De novo* Alu insertion analysis

HeLa-HA cells were seeded in 6-well plates at 1.5 × 10^5^ cells per well in MEM culture medium. Approximately 24 hours later (day 0), 1.0 µg of either cepB-Alu-gfp(427)-O2F3RT(-) or cepB-Alu-gfp(427)-O2F3 plasmid was mixed with 3.0 µL of FuGENE HD and 100 µL of Opti-MEM. After a 10-minute incubation at room temperature, the mixtures were added to the cells for transfection. On day 1, the medium was replaced with fresh MEM. Selection was initiated on day 2 with blasticidin (10 µg/mL), and on day 10, cells were harvested and washed with 1× PBS. After removing the supernatant, the cells were resuspended in 1× PBS, and an EGFP-positive cell population was sorted using a BD FACSAria III.

Recovery of insertion sequences was performed generally according to Richardson et al. (2014), with minor modifications as described below. To isolate single colonies, sorted cells were diluted and seeded onto a 10-cm dish. After approximately two weeks of culture, single colonies were embedded in 1× PBS containing SeaPlaque GTG Agarose (Lonza), harvested, and re-seeded into a 6-well plate. Cells were cultured until confluent, then harvested. Genomic DNA was extracted using the Wizard Genomic DNA Purification Kit (Promega) following the manufacturer’s protocol. Eight micrograms of genomic DNA was digested overnight at 37°C with SspI-HF, NdeI, or XbaI (New England Biolabs). The enzymes were subsequently inactivated by heating at 65°C for 30 minutes. The resulting genomic DNA fragments were self-ligated to form circular DNA in a 500 µL reaction containing 3,200 U of T4 DNA Ligase (New England Biolabs) at 16°C for 24 hours. To purify the circularized DNA, 50 µL of 3 M sodium acetate (pH 5.2) and 500 µL of isopropanol were added to the reaction, and the mixture was centrifuged at ∼20,000 × g for 15 minutes at 4°C. After removing the supernatant, the pellet was washed with 200 µL of 70% (v/v) ethanol and centrifuged again. The supernatant was discarded, and the DNA pellet was air-dried and dissolved in Buffer EB (QIAGEN).

To analyze the genomic sequences flanking the *de novo* Alu insertions, a two-step nested inverse PCR was performed using primers specific to the *EGFP* sequence. The PCR was carried out with SeqAmp DNA Polymerase (TaKaRa Bio), the Expand Long Template PCR System (Roche), or Q5 Hot Start High-Fidelity 2× Master Mix (New England Biolabs).

For reactions with SeqAmp DNA Polymerase, the first-round PCR was performed in a 25 µL mixture containing 12.5 µL of 2× SeqAmp PCR Buffer (TaKaRa Bio), 0.5 µL of SeqAmp DNA Polymerase, 2.0 µL of circularized genomic DNA, and 0.2 µM each of SMA045 and SMA046. The PCR program consisted of an initial denaturation at 94°C for 1 minute; 30 cycles of denaturation at 98°C for 10 seconds and annealing/extension at 68°C for 15 minutes; and a final extension at 68°C for 30 minutes. For the second-round PCR, 2.0 µL of the first-round product served as the template with primers SMA047 and SMA048 under identical cycling conditions. For reactions with the Expand Long Template PCR System, the first-round PCR was performed in a 20 µL mixture containing 2.0 µL of 10× Buffer, 5.0 µL of 2 mM dNTPs, 0.2 µL of Enzyme Mix, 2.0 µL of circularized genomic DNA, and 0.3 µM each of SMA045 and SMA046. The thermal cycling conditions consisted of an initial denaturation at 95°C for 2 minutes; 30 cycles of denaturation at 94°C for 10 seconds, annealing at 65°C for 30 seconds, and extension at 68°C for 15 minutes; and a final extension at 68°C for 30 minutes. The second-round PCR used 2.0 µL of the first-round product as the template with SMA047 and SMA048 under the same reaction and cycling conditions. For reactions with the Q5 Hot Start High-Fidelity 2X Master Mix, the first-round PCR was performed in a 25 µL mixture containing 12.5 µL of Q5 Hot Start High-Fidelity 2× Master Mix, 2.0 µL of circularized genomic DNA, and 0.5 µM each of SMA045 and SMA046. The thermal cycling conditions consisted of an initial denaturation at 98°C for 30 seconds; 35 cycles of denaturation at 98°C for 10 seconds, annealing at 72°C for 30 seconds, and extension at 72°C for 5 minutes; and a final extension at 72°C for 2 minutes. The second-round PCR used 2.0 µL of the first-round product as the template with SMA047 and SMA048 under the same reaction and cycling conditions.

Following PCR, the amplified products were separated by agarose gel electrophoresis. DNA fragments were excised from the gel and purified using a Gel Extraction Kit (QIAGEN) according to the manufacturer’s protocol. The purified PCR products were cloned into the pCR4 Blunt-TOPO vector (Thermo Fisher Scientific), pCR-Blunt II-TOPO vector (Thermo Fisher Scientific), or pGEM-T Easy vector (Promega) and subjected to Sanger sequencing. *De novo* Alu insertion sites were identified using the Human BLAT Search tool on the UCSC Genome Browser with the GRCh38/hg38 human genome assembly; gene annotations were based on NCBI RefSeq. To confirm the Alu-*EGFP*^Tet^ insertion at the identified loci, the surrounding genomic regions were amplified by PCR. PCR reactions were carried out in 25 µL containing 0.3 µg of genomic DNA, 12.5 µL of 2× SeqAmp PCR Buffer, 0.5 µL of SeqAmp DNA Polymerase, and 0.8 µM of each primer in the corresponding primer pair: SMA074/SMA075 (#1), SMA055/SMA056 (#2), SMA315/SMA316 (#3), SMA076/SMA077 (#4), or SMA317/SMA318 (#5). The PCR program consisted of an initial denaturation at 94°C for 1 minute; 30 cycles of denaturation at 98°C for 10 seconds and annealing/extension at 68°C for 2 minutes; and a final extension at 68°C for 3 minutes. PCR products were then resolved on a 1.5% (w/v) agarose gel, stained with GelRed, and visualized under UV light using an E-box CX5 imager. To confirm the absence of the self-splicing intron and estimate the 3′ poly(A)-tract length, locus-specific PCR products containing the insertion were cloned into pCR4 Blunt-TOPO, and plasmid DNA from two independent bacterial colonies per insertion was subjected to Sanger sequencing. Because of the instability of long homopolymeric sequences during PCR and cloning, poly(A)-tract lengths are described as approximate values.

### Microscopy analysis of EGFP-positive cells

As described above, HeLa-HA cells were transfected with cepB-Alu-gfp(427)-O2F3RT(-) or cepB-Alu-gfp(427)-O2F3. On day 8 after transfection, cells were harvested and washed with 1× PBS. After removing the supernatant, cells were resuspended in 1× PBS containing 2% (v/v) FBS and 0.5 mM EDTA. Fifty thousand cells from each of the RT-deficient, EGFP-negative, and EGFP-positive populations were sorted using a BD FACSAria IIu. Sorted cells were cultured and imaged on day 14 after transfection using a BZ-X810 all-in-one fluorescence microscope (KEYENCE) with a Plan Fluorite 20× LD PH (BZ-PF20LP) objective lens. EGFP fluorescence was observed using the BZ-X GFP filter, and images were viewed using BZ-X800 Viewer (version 1.3.0.1). Background correction was performed using BZ-X800 Analyzer (version 1.1.2.4), and all images were processed under identical conditions.

### Western blotting

HeLa-HA cells were seeded into 6-well culture plates with 2 mL of MEM culture medium at 2.0 × 10^5^ cells per well. Twenty-four hours later, siRNAs targeting the indicated genes were introduced into the cells as described above. On day 3 after siRNA treatment, cells were washed with 1× PBS, treated with 0.25% Trypsin-EDTA, and resuspended in MEM culture medium. After three washes with 1× PBS, cells were quickly frozen in liquid nitrogen and stored at -80°C. To extract proteins, 40–60 µL of Radio-ImmunoPrecipitation Assay (RIPA) buffer (10 mM Tris-HCl [pH 7.5], 1 mM EDTA [Nacalai Tesque], 1% [v/v] Triton X-100 [Nacalai Tesque], 0.1% [w/v] sodium deoxycholate [Sigma-Aldrich], 0.1% [w/v] SDS [Nacalai Tesque], 140 mM NaCl [Nacalai Tesque], and 1× cOmplete EDTA-free protease inhibitor cocktail [Roche]) was added to each cell pellet, and the suspension was incubated at 4°C for 30 minutes. Then, the resulting cell lysates were centrifuged at ∼13,000 × g for 5 minutes at 4°C, and the supernatant was collected. Protein concentrations were measured using the Bio-Rad Protein Assay Kit with an iMark microplate reader (Bio-Rad). Samples were adjusted to 4 µg/µL with 3× SDS-PAGE sample buffer (0.188 M Tris-HCl [pH 6.8], 30% [v/v] glycerol [Nacalai Tesque], 6.0% [w/v] SDS, 0.3 M DTT [Nacalai Tesque], and 0.02% [w/v] bromophenol blue [Nacalai Tesque]) and boiled at 105°C for 4 minutes. Then, 10‒20 µg of protein was separated by SDS-PAGE and transferred to an Immobilon-FL PVDF membrane (Merck Millipore) using semi-dry buffer (24 mM Tris, 0.1% [w/v] SDS, 192 mM glycine [Nacalai Tesque], 20% [v/v] ethanol [Nacalai Tesque]) in a Trans-Blot SD Semi-Dry Transfer Cell (Bio-Rad) at 10 V for 1 hour (SRP9/14) or in 10 mM CAPS buffer (3-[cyclohexylamino]-1-propanesulfonic acid [Nacalai Tesque], pH 11.0) in a Mini Trans-Blot Electrophoretic Transfer Cell (Bio-Rad) at 50 V for 12 hours at 4°C (MOV10). After transfer, membranes used for SRP9, SRP14, and α-tubulin detection were blocked with 3% (w/v) skim milk (MEGMILK SNOW BRAND) in 1× TNT buffer (20 mM Tris-HCl [pH 7.5], 140 mM NaCl, and 0.05% [v/v] Tween 20), whereas those used for MOV10 and GAPDH detection were blocked with Intercept Blocking Buffer (LI-COR Biosciences). After blocking for 1 hour at room temperature, membranes were incubated overnight at 4°C with primary antibodies diluted in 1× TNT buffer (SRP9/14) or Intercept Blocking Buffer (MOV10). The next day, membranes were washed four times with 1× TNT buffer and incubated overnight at 4°C with secondary antibodies diluted in 1× TNT buffer containing 0.01% (w/v) SDS. Finally, membranes were washed four times with 1× TNT buffer, followed by one wash with 1× PBS. Signals were detected using an Odyssey imaging system with LI-COR Acquisition Software (version 1.2) and analyzed using Empiria Studio Software (version 2.3).

### Plasmids used in this study

All plasmids used for cell transfection were purified using the Plasmid Midiprep Kit (Qiagen), the GenElute HP Plasmid Miniprep Kit (Sigma-Aldrich), or the PureLink HiPure Plasmid Miniprep Kit (Thermo Fisher Scientific). Unless otherwise indicated, L1-derived constructs were based on the human L1.3 element (GenBank accession no. L19088). The LRE3 element used in this study was originally described in Brouha et al. (2002). Amino acid positions in ORF2p are numbered from the first methionine of the ORF2 sequence.

pKO17_pBSAluegfp(493): This reporter plasmid uses a pBlueScript SK(-) backbone and contains the AluY element used previously (Dewannieux et al., 2003; Doucet et al., 2015; Moldovan et al., 2024, 2025). An *EGFP* reporter cassette was inserted between the 3′ end of the AluY sequence and the encoded 44-adenine poly(A) tract, in the antisense orientation relative to Alu transcription. The *EGFP* coding sequence is interrupted by a *Tetrahymena* group I self-splicing intron (Dewannieux et al., 2003; Esnault et al., 2002). The *EGFP* exon sequences flanking the insertion site and the intron IGS were modified to preserve P1 and P10 base pairing without altering the encoded EGFP amino acid sequence; the intron was oriented in the same direction as Alu transcription. The intron is located 493 bp downstream of the first nucleotide of the *EGFP* start codon. *EGFP* expression is driven by an SV40 promoter. A 7SL enhancer is located upstream of Alu to enhance Pol III transcription. Two TTTT motifs downstream of the encoded poly(A) tract serve as Pol III termination signals.

pKO18_pBSAluegfp(393): This plasmid is identical to pKO17_pBSAluegfp(493) except that the self-splicing intron is located 393 bp downstream of the *EGFP* start codon.

pKO19_pBSAluegfp(427): This plasmid is identical to pKO17_pBSAluegfp(493), except that the self-splicing intron is located 427 bp downstream of the *EGFP* start codon.

pSMA18_pBSAluegfp(-intron): This plasmid is identical to pKO17_pBSAluegfp(493) except it lacks the self-splicing intron.

pSMA72_pBSAluEFSegfp(427): This reporter plasmid is a derivative of pKO19_pBSAluegfp(427), generated by replacing the SV40 promoter with an EFS promoter.

pTMO2F3: This plasmid, described previously (Doucet et al., 2015; Miyoshi et al., 2019), expresses monocistronic ORF2p derived from L1.3, fused with a C-terminal 3×FLAG tag. Expression is driven by a CMV promoter and the native L1 5′ UTR. The plasmid contains a hygromycin-resistance gene for selection and lacks a retrotransposition reporter cassette.

pTMO2F3D702A: This plasmid is a derivative of pTMO2F3 and has been previously described (Doucet et al., 2015; Miyoshi et al., 2019). It is identical to pTMO2F3 except for a D702A point mutation in the reverse transcriptase (RT) domain of ORF2p (Wei et al., 2001), which inactivates the RT activity. The plasmid was used as a negative control for retrotransposition.

pSMA92_pEF1O2F3_B: This plasmid is a derivative of pTMO2F3_B, a version of pTMO2F3 in which the hygromycin-resistance gene was replaced with a blasticidin-resistance gene. It expresses a monocistronic ORF2p-3×FLAG derived from L1.3 under the EF1α promoter rather than the CMV promoter, and retains the native L1 5′ UTR. The blasticidin-resistance gene is driven by the SV40 promoter.

pSMA93_pEF1O2F3D702A_B: This plasmid is identical to pSMA92_pEF1O2F3_B but contains the D702A mutation in ORF2p.

pSMA186_pEF1O2F3LRE3_B: This plasmid is a derivative of pSMA92_pEF1O2F3_B, in which the L1.3-derived ORF2 coding sequence was replaced with the LRE3-derived ORF2 coding sequence (Brouha et al., 2002; Farley et al., 2004). ORF2p expression is driven by an EF1α promoter and the native L1.3 5′ UTR. The plasmid also contains a blasticidin-resistance gene for selection.

pTM541_cepB-Alu-gfp(427)-L1F3: This plasmid was generated by inserting the Alu-*EGFP*^Tet^ construct derived from pKO19_pBSAluegfp(427) into pTMF3_B at the NruI site. In pTMF3_B, the hygromycin-resistance gene was replaced with a blasticidin-resistance gene, as in pTMO2F3_B. pTMF3_B retains the CMV promoter. pTM541 expresses Alu RNA and the full-length L1. pTMF3, described previously (Miyoshi et al., 2019), encodes a full-length L1.3 with a T7 epitope-tagged ORF1p and a 3×FLAG-tagged ORF2p.

pTM542_cepB-Alu-gfp(493)-L1F3: This plasmid was generated by inserting the Alu-*EGFP*^Tet^ construct derived from pKO17_pBSAluegfp(493) into pTMF3_B at the NruI site.

pTM545_cepB-Alu-gfp(393)-L1F3: This plasmid was generated by inserting the Alu-*EGFP*^Tet^ construct derived from pKO18_pBSAluegfp(393) into pTMF3_B at the NruI site.

pTM543_cepB-Alu-gfp(427)-O2F3: This plasmid was generated by inserting the Alu-*EGFP*^Tet^ construct derived from pKO19_pBSAluegfp(427) into pTMO2F3_B at the NruI site. It expresses Alu RNA and ORF2p fused with three copies of a FLAG tag at the C-terminus.

pTM544_cepB-Alu-gfp(493)-O2F3: This plasmid was generated by inserting the Alu-*EGFP*^Tet^ construct derived from pKO17_pBSAluegfp(493) into pTMO2F3_B at the NruI site.

pTM546_cepB-Alu-gfp(393)-O2F3: This plasmid was generated by inserting the Alu-*EGFP*^Tet^ construct derived from pKO18_pBSAluegfp(393) into pTMO2F3_B at the NruI site.

pTM549_cepB-Alu-gfp(427)-O2F3RT(-): This plasmid is identical to pTM543_cepB-Alu-gfp(427)-O2F3, except that it carries the D702A point mutation in the RT domain of ORF2p. It was used as a negative control for retrotransposition.

cepB-gfp-L1.3: This plasmid, previously described (Miyoshi et al., 2019), contains a full-length human L1.3 element with an *EGFP* retrotransposition indicator cassette (*mEGFPI*) inserted

into its 3′ UTR (Ostertag et al., 2000). The entire construct is carried in a modified pCEP4 vector that includes a blasticidin resistance gene for selection. Expression of the engineered L1 is driven by its native 5′ UTR.

pSMA177_pBSAluYa5egfp(427): This reporter plasmid is a derivative of pKO19_pBSAluegfp(427) generated by replacing the AluY sequence with the AluYa5 consensus sequence (Jurka, 2000).

pSMA181_pBSAluYb8egfp(427): This reporter plasmid is a derivative of pKO19_pBSAluegfp(427) generated by replacing the AluY sequence with the AluYb8 consensus sequence (Jurka, 2000).

pSMA41_pBSB2egfp(427): This reporter plasmid is a derivative of pKO19_pBSAluegfp(427), generated by replacing the AluY sequence with a mouse B2 element. The B2 sequence is identical to that used previously in a neomycin resistance-based reporter assay (Dewannieux & Heidmann, 2005a).

pSMA40_pBSAluecfp(427): This reporter plasmid is a derivative of pKO19_pBSAluegfp(427) generated by introducing four amino acid substitutions (Y66W, N146I, M153T, and V163A) into the *EGFP* sequence.

pSMA44_pBSAlucerulean(427): This reporter plasmid is a derivative of pSMA40_pBSAluecfp(427) generated by introducing three amino acid substitutions (S72A, Y145A, H148D) into the *ECFP* sequence.

pSMA77_pBSAlumTurquoise2(427): This reporter plasmid is a derivative of pSMA44_pBSAlucerulean(427) generated by introducing five amino acid substitutions (T65S, A145Y, I146F, S175G, A206K) into the *Cerulean* sequence.

pSMA78_pBSAluEFSmTurquoise2(427): This reporter plasmid is a derivative of pSMA77_pBSAlumTurquoise2(427), generated by replacing the SV40 promoter with the EFS promoter.

pBSmock: This plasmid is the pBlueScript SK(-) backbone used as a mock control.

pSMA116_pBSAluΔA44EFSmTurquoise2(427): This reporter plasmid is a derivative of pSMA78_pBSAluEFSmTurquoise2(427) generated by deleting the 44-adenine poly(A) tract at the 3′ end of the Alu sequence.

pTM587_cep99-gfp-EF1LRE3: This plasmid is a derivative of cep99-gfp-L1, as described previously (Miyoshi et al., 2019; Ostertag et al., 2000). It contains the full-length LRE3 element originally described by Brouha et al. (2002) instead of L1.3, uses the EF1α promoter for L1 transcription, carries the *mEGFPI* retrotransposition indicator cassette, and served as the wild-type L1 reporter in the dual-color assay. The plasmid also contains a puromycin-resistance gene for selection.

pTM608_cep99-gfp-EF1LRE3RR261_262AA: This plasmid is a derivative of pTM587_cep99-gfp-EF1LRE3 that encodes the ORF1p R261A/R262A substitutions.

### Primers used in this study

KO69: 5’-CGACGAGGCCCAGAGCA-3’

KO70: 5’-GTCCACGTCACACTTCATGATGGA-3’

KO87: 5’-CACGAACTCCAGCAGGACCA-3’

KO88: 5’-GGCCACAAGTTCAGCGTGTC-3’

SMA045: 5’-CGCGGGTCTTGTAGTTGCCGTCG-3’

SMA046: 5’-CACCCAGTCCGCCCTGAGCAAAG-3’

SMA047: 5’-AGAAGATGGTGCGCTCCTGGACG-3’

SMA048: 5’-CGAGAAGCGCGATCACATGGTCC-3’

SMA055: 5’-AGCTAAGAAGGGAAGCAAAGGA-3’

SMA056: 5’-TGGGTAGAGTGATACTTCCAGC-3’

SMA074: 5’-GCTTCCAGTGGCACTAGGAT-3’

SMA075: 5’-TCGTGACTGCTGTATTCACTGT-3’

SMA076: 5’-GCTCACACAAATTAAAAGCAGCA-3’

SMA077: 5’-TGTCACCAGCAGTAAATGAACA-3’

SMA315: 5’-CACCCTTGAGTGGCTCATGT-3’

SMA316: 5’-TCCCTTCCAAGATAGGTGACA-3’

SMA317: 5’-AGTGTGTGGGTCTAGTGGAGA-3’

SMA318: 5’-TGCATGTGGATGTACCCTGTA-3’

oligo (dT): 5’-TTTTTTTTTTTTTTTTTTTTVN-3’

### Antibodies used in this study

Rabbit polyclonal anti-SRP9 antibody (1:1,000; Proteintech, 11195-1-AP)

Rabbit polyclonal anti-SRP14 antibody (1:1,000; Proteintech, 11528-1-AP)

Mouse monoclonal anti-α-Tubulin antibody (1:1,000; Sigma-Aldrich, T9026-2ML)

Rabbit polyclonal anti-MOV10 antibody (1:1,000; Proteintech, 10370-1-AP)

Mouse monoclonal anti-GAPDH antibody, clone 6C5 (1:2,000; Sigma-Aldrich, MAB374)

IRDye 680RD anti-mouse IgG (1:10,000; LI-COR Biosciences, 925-68072)

IRDye 680RD anti-rabbit IgG (1:10,000; LI-COR Biosciences, 926-68071)

IRDye 800CW anti-mouse IgG (1:10,000; LI-COR Biosciences, 926-32212)

IRDye 800CW anti-rabbit IgG (1:10,000; LI-COR Biosciences, 926-32213)

### Statistical analysis

Quantitative data are presented as the mean ± SEM from three independent experiments unless otherwise stated. For quantitative flow-cytometry assays, each condition was analyzed in technical duplicate, and the two technical replicates were averaged to obtain one value per biological replicate for the quantitative graphs and statistical analyses.

Two-sided Welch’s t-tests were used for the Alu–B2 comparison in Fig. 4e, the intact-Alu comparison between wild-type and ORF1p-mutant L1 in Fig. 5f, and the *EGFP*^Tet^–EFS-*EGFP*^Tet^ comparison in Fig. 5c. The three comparisons between corresponding monocistronic ORF2p and full-length L1 conditions in Fig. 2d and the two L1.3–LRE3 ORF2p comparisons in Fig. 4f were performed using two-sided Welch’s t-tests, with Holm adjustment applied separately within each figure panel.

Welch’s one-way ANOVA followed by Games–Howell tests was used for the mock, intact Alu and poly(A)-deficient Alu conditions under wild-type L1 in Fig. 5g, the three active Alu-subfamily reporters in Fig. 4e, the four cyan reporter variants in Fig. 5b, and the Alu EGFP-positive fractions in Supplementary Fig. 3c.

Relative reporter activities in Fig. 4c were compared with a theoretical mean of 1.0 using two-sided one-sample t-tests; the SRP9 and SRP14 knockdown comparisons were adjusted using the Holm method. For the HeLa-HA time course data in Fig. 4g, the L1.3- and LRE3-derived ORF2p conditions were compared at each time point using two-sided Welch’s t-tests with Holm adjustment across the seven time points.

For the observed-versus-expected analyses, event counts from two technical replicates of 500,000 singlet events each were pooled within each biological replicate, yielding counts per 10^6^ singlet events. Expected dual-positive counts under independence were calculated as (mTurquoise2-positive events × EGFP-positive events) / total singlet events. Observed and expected counts are shown in Fig. 5i and Supplementary Fig. 3e; observed-to-expected ratios are provided in Supplementary Table 1.

Statistical analyses were performed using R (version 4.3.1; R Foundation for Statistical Computing). P < 0.05 was considered statistically significant.

## Data availability

The source data used for the quantitative analyses are provided in Supplementary Tables 1 and 2. Additional data and plasmids generated in this study are available from the corresponding author upon reasonable request.

## Code availability

No custom software was used in this study.

## Acknowledgements

We thank J. V. Moran, A. M. Roy-Engel, and T. Heidmann for valuable reagents; J. Suzuki, K. Aoki, and T. Uemura, all Ishikawa lab members at Kyoto University, and all lab members in the RIKEN IMS Laboratory for Retrotransposon Dynamics for helpful discussions. S.A. and K.N. were supported by the RIKEN Junior Research Associate (JRA) and RIKEN Student Researcher (RSR) programs. T.M. was supported by JSPS KAKENHI (Grant Numbers 21K19219, 23K23863, 26K01990, 26K23183, and 26H01551), JST PRESTO (Grant Number JPMJPR2289), and research grants from the Takeda Science Foundation, Astellas Foundation for Research on Metabolic Disorders, Nagase Science Technology Foundation, and G-7 Scholarship Foundation. We also thank Takako Tsuda, Yoko Hirata, and Haruno Suzuki for their excellent secretarial support.

## Author information

### Contributions

S.A., K.O., and T.M. conceived and designed the experiments and analyzed the data. S.A., K.O., K.N., and T.M. performed experiments. S.A. and T.M. wrote and edited the manuscript with input from all authors. All authors discussed the results and approved the final manuscript.

### Corresponding author

Please address correspondence to Tomoichiro Miyoshi.

## Supplementary information

Supplementary Figures 1-3, Supplementary Tables 1-2, and Supplementary Data 1.

