## Supplementary material for "Dual-color fluorescent reporters resolve Alu and LINE-1 retrotransposition in single cells": Table 1

Contents

Supplementary Figure 1

Supplementary Figure 2

Supplementary Figure 3

Supplementary Table 1 legend

Supplementary Table 2 legend

Supplementary Data 1

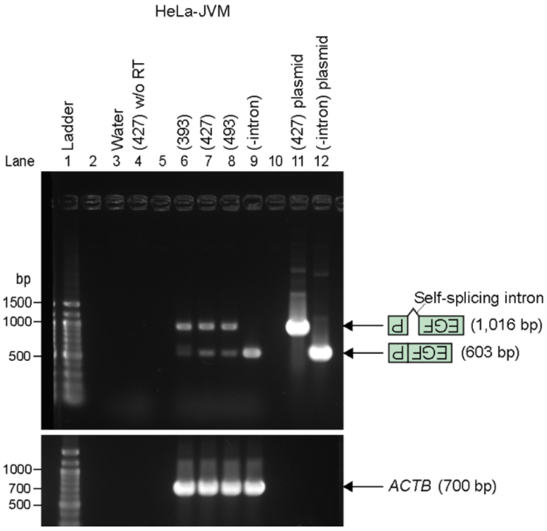


**Supplementary Figure 1. Detection of intron-free Alu-*EGFP*$^{Tet}$ reporter transcripts in HeLa-JVM cells**.

RT-PCR analysis of RNA from transfected HeLa-JVM cells. The expected sizes of the cDNA products containing or lacking the self-splicing intron are ~1.0 kb and ~0.6 kb, respectively. *ACTB*, used as a reverse transcription control, was detected at ~0.7 kb. Lanes "(393)," "(427)," and "(493)" correspond to pBSAluegfp(393), pBSAluegfp(427), and pBSAluegfp(493), respectively. Negative controls include water and RNA from pBSAluegfp(427)-transfected cells without reverse transcription. PCR-amplified DNA from the intron-containing pBSAluegfp(427) plasmid and the intronless pBSAluegfp(-intron) plasmid were included as controls.

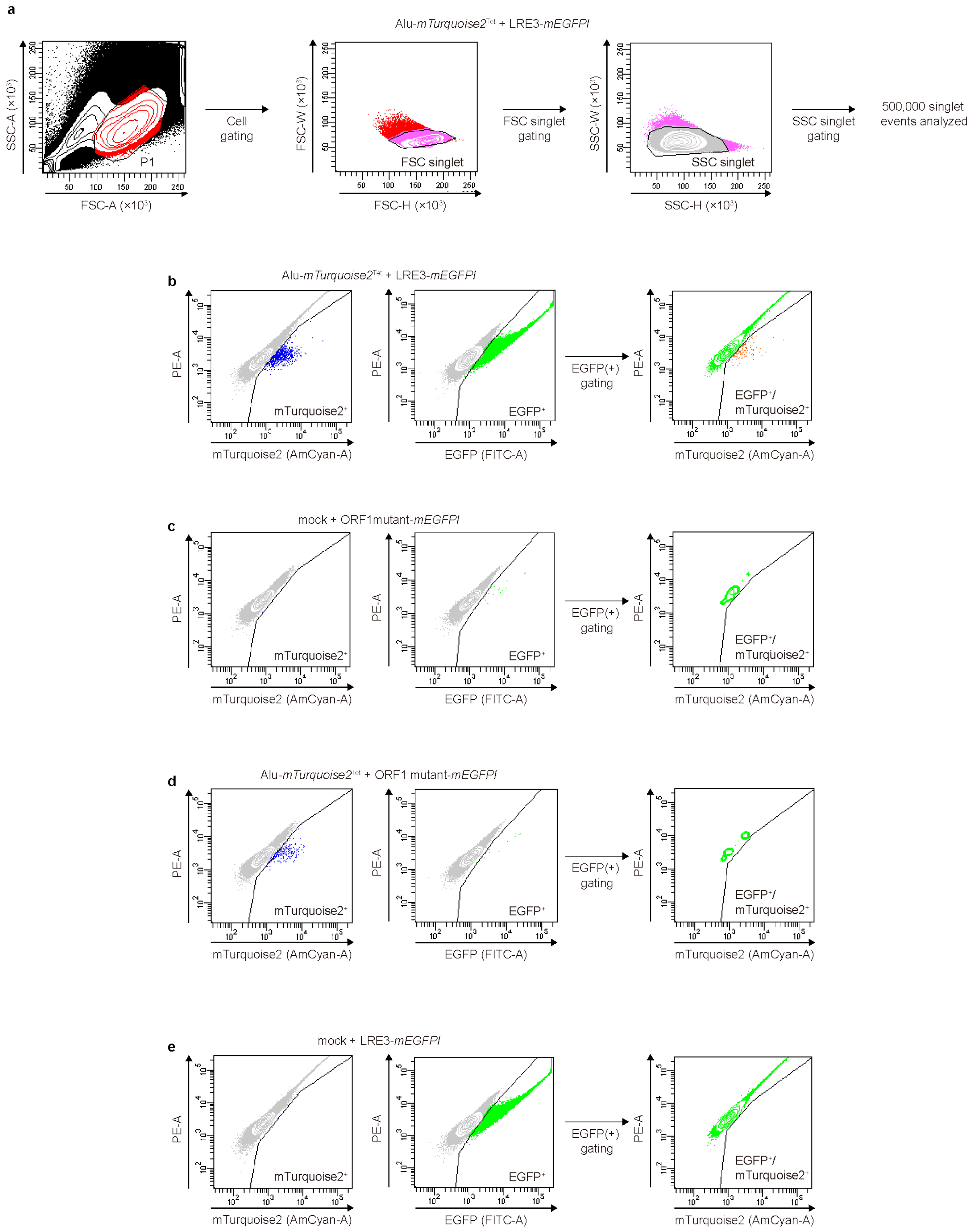


**Supplementary Figure 2. Gating strategy for a dual-color retrotransposition assay.**

**a**, Sequential gating strategy used for the dual-color flow-cytometric analysis. Cells were first gated on an SSC-A versus FSC-A contour plot (P1), followed by sequential singlet gating using FSC-W versus FSC-H and SSC-W versus SSC-H. For the representative analysis shown, 500,000 singlet events were retained for fluorescence analysis. **b**, Fluorescence gating of cells co-transfected with intact Alu-EFS-*mTurquoise2*$^{\mathrm{Tet}}$ and wild-type LRE3-

*mEGFPI*. mTurquoise2-positive and EGFP-positive populations were identified using PE-A versus AmCyan-A and PE-A versus FITC-A plots, respectively. EGFP-positive events were then gated and analyzed for mTurquoise2 fluorescence to identify EGFP/mTurquoise2 dual-positive events. **c–e**, The same gating strategy was applied to corresponding control conditions used to define the fluorescence gates: mock plus ORF1p-mutant LRE3-*mEGFPI*, representing the dual-negative control (**c**); intact Alu-EFS-*mTurquoise2*$^{Tet}$ plus ORF1p-mutant LRE3-*mEGFPI*, representing the mTurquoise2-active/EGFP-background control (**d**); and mock plus wild-type LRE3-*mEGFPI*, representing the EGFP-active/mTurquoise2-negative control (**e**). The same fluorescence gates were applied across all conditions. Gray events indicate the parent population; blue, green and orange events indicate mTurquoise2-positive, EGFP-positive and EGFP/mTurquoise2 dual-positive populations, respectively. PE-A denotes fluorescence collected through the 585/42-nm band-pass filter after 488-nm excitation.

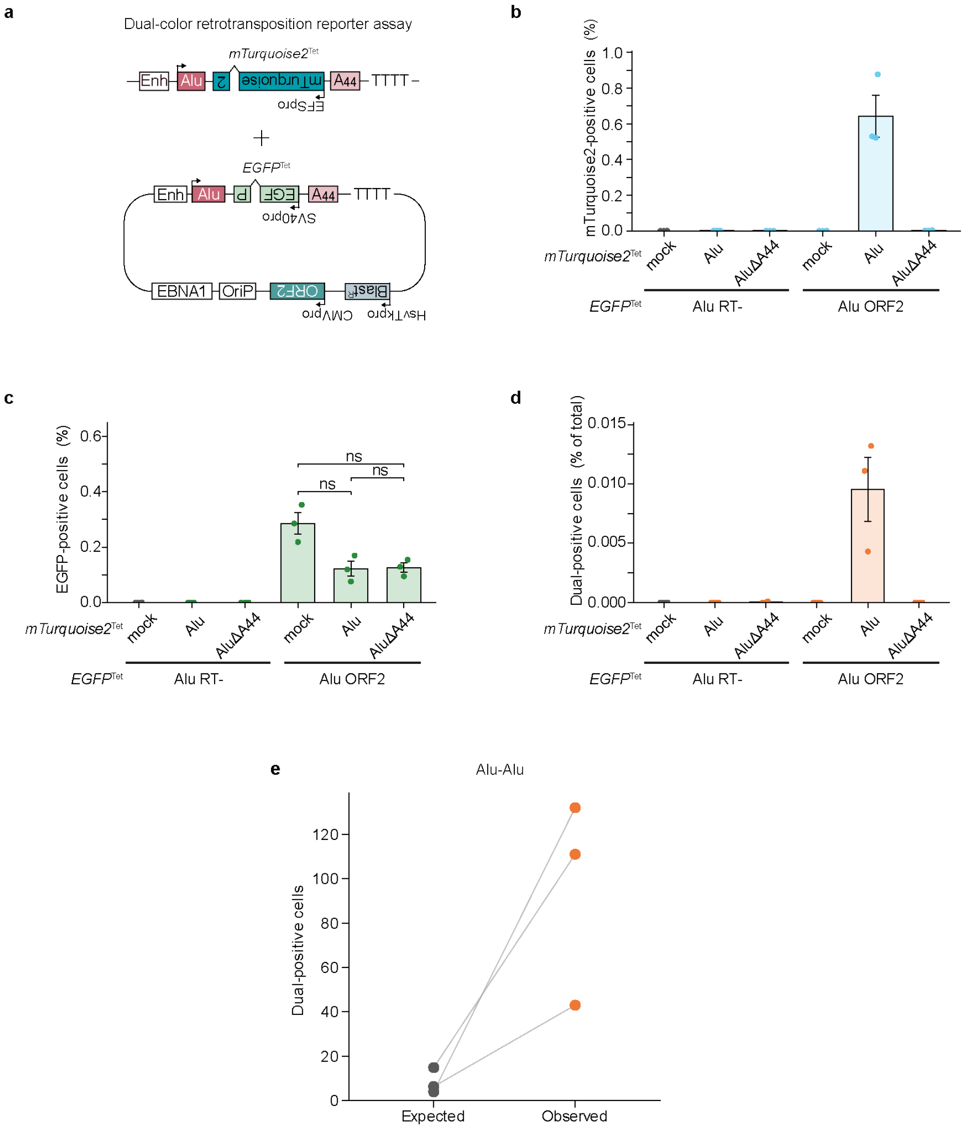


**Supplementary Figure 3. Dual-color analysis of two Alu reporters.**

**a**, Schematic of the Alu–Alu dual-color assay. Alu-*EGFP*$^{\text{Tet}}$ and Alu-EFS-*mTurquoise2*$^{\text{Tet}}$ were co-transfected into HeLa-HA cells, and retrotransposition was driven by monocistronic wild-type or RT-deficient ORF2p. **b**, mTurquoise2-positive fractions under the indicated ORF2p and cyan-reporter conditions. **c**, EGFP-positive fractions derived from Alu-*EGFP*$^{\text{Tet}}$ in the wild-type ORF2p background after co-transfection with mock, intact cyan Alu or poly(A)-deficient cyan Alu (ΔA44). The same samples as in b were analyzed. **d**, EGFP/mTurquoise2 dual-positive fractions in the same samples as in b. Dual-positive fractions were calculated relative to the total number of analyzed singlet events. **e**, Observed dual-positive event counts compared with counts expected under independence in the intact Alu–Alu condition. Event counts from two 500,000-singlet technical replicates were pooled within each biological replicate and are shown per 1,000,000 singlet events; lines connect values from the same biological replicate. Expected counts were calculated as described in Supplementary Table 1. Data in **b–d** are mean ± SEM from three independent experiments. The three conditions in **c**

were compared using Welch's one-way ANOVA followed by Games–Howell tests. ns, not significant.

**Supplementary Table 1. Observed and expected dual-positive events in the Alu–L1 and Alu–Alu dual-color assays.**

For each biological replicate, the table lists the total singlet count pooled from two 500,000-event technical replicates, mTurquoise2-positive and EGFP-positive event counts and frequencies, observed dual-positive event counts and frequencies, expected dual-positive counts and frequencies under independence, and the observed-to-expected ratio. Expected dual-positive counts were calculated as (mTurquoise2-positive events × EGFP-positive events) / total singlet events.

**Supplementary Table 2. Plasmid combinations, experimental conditions and raw flow-cytometry data supporting the quantitative analyses.**

The first worksheet lists, for each figure panel, the cell line, reporter and driver plasmids, selection condition, and detection channel. Subsequent worksheets provide raw event counts and reporter-positive frequencies for all quantitative flow-cytometry analyses shown in the main and supplementary figures. Each raw-data worksheet lists the experimental condition, biological replicate, technical replicate, total number of analyzed events, reporter-positive event counts, corresponding frequencies, and, where applicable, statistical results. Technical replicates were analyzed separately and averaged to obtain one value per biological replicate for quantitative graphs and statistical analyses, except for the observed-versus-expected analyses described in Supplementary Table 1. For the dual-color assays, the worksheets also include dual-positive event counts for matched control conditions in which only one reporter was detectably active. These counts provide empirical estimates of dual-gated background, including reciprocal cross-gating, autofluorescence, and gate-boundary events.

**Supplementary Data 1. Sequences of recovered Alu-*EGFP*$^{Tet}$ insertions.**

For each insertion, bold text denotes the inserted Alu-*EGFP*$^{Tet}$ sequence, underlined text denotes the target-site duplication (TSD), and unformatted text denotes flanking genomic DNA. Extra nucleotides that could not be assigned to either the reporter or the flanking genomic sequence are shown at the corresponding insertion junctions and are summarized in Table 1. The 3′ poly(A) tracts are represented as (A)n; approximate tract lengths are provided in Table 1.

#1
GCTTCCAGTGGCACTAGGATGAGTACAATTTCAGTTTCATTGCATAGAGAAGCATTAAAACA
TTTTTATTCGATACTATAATTGGAAATGAGTTTCTTGTGATAGTGCTTGTGACAGTAACATTCA
AAGTCTACATTTAACCACAGTAATTGCGATATGTGTAAGAATGAAAAATCCT

A**GGCCGGGCGCGGTGGCTCACGCCTGTAATCCCAGCACTTTGGGAGGCCGAGGCGGGC**
**GGATCACGAGGTCAGGAGATCGAGACCATCCTGGCTAACACGGTGAAACCCCGTCTCTA**
**CTAAAAAAAAAATACAAAAAAATTAGCCGGGCGTGGTAGCGGGCGCCTGTAGTCCCAGC**
**TACTCGGGAGGCTGAGGCAGGGGAATGGCGTGAACCCGGGAGGCGGAGCTTGCAGTGA**
**GCCGAGATCCCGCCACTGCACTCCAGCCTGGGCGACAGAGCGAGACTCCGTCTCAAATC**
**CCCCTACTTGTACAGCTCGTCCATGCCGAGAGTGATCCCGGCGGCGGTCACGAACTCCA**
**GCAGGACCATGTGATCGCGCTTCTCGTTGGGGTCTTTGCTCAGGGCGGACTGGGTGCTCA**
**GGTAGTGGTTGTCGGGCAGCAGCACGGGGCCGTCGCCGATGGGGGTGTTCTGCTGGTAG**
**TGGTCGGCGAGCTGCACGCTGCCGTCCTCGATGTTGTGGCGGATCTTGAAGTTCACCTTG**
**ATGCCGTTCTTCTGCTTGTCGGCCATGATATAGACGTTGTGGCTGTTGTAGTTGTACTCAA**
**GCTTGTGCCCCAGGATGTTGCCGTCCTCCTTGAAGTCGATGCCCTTCAGCTCGATGCGGT**
**TCACCAGGGTGTCGCCCTCGAACTTCACCTCGGCGCGGGTCTTGTAGTTGCCGTCGTCCT**
**TGAAGAAGATGGTGCGCTCCTGGACGTAGCCTTCGGGCATGGCGGACTTGAAGAAGTCG**
**TGCTGCTTCATGTGGTCGGGGTAGCGGCTGAAGCACTGCACGCCGTAGGTCAGGGTGGT**
**CACGAGGGTGGGCCAGGGCACGGGCAGCTTGCCGGTGGTGCAGATGAACTTCAGGGTC**
**AGCTTGCCGTAGGTGGCATCGCCCTCGCCCTCGCCGGACACGCTGAACTTGTGGCCGTTT**
**ACGTCGCCGTCCAGCTCGACCAGGATGGGCACCACCCCGGTGAACAGCTCCTCGCCCTT**
**GCTCACCATATTGGCTCGACCTCCAAAAAAGCCTCCTCACTACTTCTGGAATAGCTCAGA**
**GGCCGAGGCGGCCTCGGCCTCTGCATAAATAAAAAAAATTAGTCAGCCATGGGGCGGAG**
**AATGGGCGGAACTGGGCGGAGTTAGGGGCGGGATGGGCGGAGTTAGGGGCGGGACTAT**
**GGTTGCTGACTAATTGAGATGCATGCTTTGCATACTTCTGCCTGCTGGGGAGCCTGGGGA**
**CTTTCCACACCTGGTTGCTGACTAATTGAGATGCATGCTTTGCATACTTCTGCCTGCTGGG**
**GAGCCTGGGGACTTTCCACACCCTAACTGACACACATTCCACAGCCGTCTC(A)n**

GAAAAATCCTCTGGTACTTTAATAGTGCAAAGACTGCAACAGTGAATACAGCAGTCACGA

#2
AGCTAAGAAGGGAAGCAAAGGACTTCTTCAAGGAGAACTGCAAACCACTGCTCAAAGAAAT
CAGAAATGGCACAAATAGAAAAACATTCCATGCTTATGAATAGGAAGAATCAATATTGTGAAA
ATGGCCATACTGTCCAAAATAATTTATAGATTCAATGCTATTCCCATTAAACTACCATTGATAT
TCTTCACAGAATTAGAAAAACCTATTTTAAAATTCATGTG

**GGCCGGGCGCGGTGGCTCACGCCTGTAATCCCAGCACTTTGGGAGGCCGAGGCGGGCG
GATCACGAGGTCAGGAGATCGAGACCATCCTGGCTAACACGGTGAAACCCCGTCTCTACT
AAAAAAAAAATACAAAAAATTAGCCGGGCGTGGTAGCGGGCGCCTGTAGTCCCAGCTAC
TCGGGAGGCTGAGGCAGGGGAATGGCGTGAACCCGGGAGGCGGAGCTTGCAGTGAGCC
GAGATCCCGCCACTGCACTCCAGCCTGGGCGACAGAGCGAGACTCCGTCTCAAATCCCC
CTACTTGTACAGCTCGTCCATGCCGAGAGTGATCCCGGCGGCGGTCACGAACTCCAGCA
GGACCATGTGATCGCGCTTCTCGTTGGGGTCTTTGCTCAGGGCGGACTGGGTGCTCAGGT
AGTGGTTGTCGGGCAGCAGCACGGGGCCGTCGCCGATGGGGGTGTTCTGCTGGTAGTGG
TCGGCGAGCTGCACGCTGCCGTCCTCGATGTTGTGGCGGATCTTGAAGTTCACCTTGATG
CCGTTCTTCTGCTTGTCGGCCATGATATAGACGTTGTGGCTGTTGTAGTTGTACTCAAGCT
TGTGCCCCAGGATGTTGCCGTCCTCCTTGAAGTCGATGCCCTTCAGCTCGATGCGGTTCA
CCAGGGTGTCGCCCTCGAACTTCACCTCGGCGCGGGTCTTGTAGTTGCCGTCGTCCTTGA
AGAAGATGGTGCGCTCCTGGACGTAGCCTTCGGGCATGGCGGACTTGAAGAAGTCGTGC
TGCTTCATGTGGTCGGGGTAGCGGCTGAAGCACTGCACGCCGTAGGTCAGGGTGGTCAC
GAGGGTGGGCCAGGGCACGGGCAGCTTGCCGGTGGTGCAGATGAACTTCAGGGTCAGCT
TGCCGTAGGTGGCATCGCCCTCGCCCTCGCCGGACACGCTGAACTTGTGGCCGTTTACGT
CGCCGTCCAGCTCGACCAGGATGGGCACCACCCCGGTGAACAGCTCCTCGCCCTTGCTC
ACCATATTGGCTCGACCTCCAAAAAAGCCTCCTCACTACTTCTGGAATAGCTCAGAGGCC
GAGGCGGCCTCGGCCTCTGCATAAATAAAAAAAATTAGTCAGCCATGGGGCGGAGAATG
GGCGGAACTGGGCGGAGTTAGGGGCGGGATGGGCGGAGTTAGGGGCGGGACTATGGTT
GCTGACTAATTGAGATGCATGCTTTGCATACTTCTGCCTGCTGGAGAGCCTGGGGACTTTC
CACACCTGGTTGCTGACTAATTGAGATGCATGCTTTGCATACTTCTGCCTGCTGGGGAGCC
TGGGGACTTTCCACACCCTAACTGACACACATTCCACAGCCGTCTC(A)n**C

AAAATTCATGTGGAACCAAAAAAGAACTCAAGTGGCCAAGACAATCCTAAGCAAAAAGAACA
AAGCTGGAAGTATCACTCTACCCA

#3

CACCCTTGAGTGGCTCATGTGAGGATACCAACCTAAATGAGTTTACCTCTTAGAGAAGATGT
CAGTGGTTGTTTTCAGTCCCATCCTCTCACTTCTCTCTGGCTTCTTTTTTAGGACGTGGAGAA
CAGCTTCTTCTTGAATGTCAATTCCCAAGTA

**GGCCGGGCGCGGTGGCTCACGCCTGTAATCCCAGCACTTTGGGAGGCCGAGGCGGGCG**
**GATCACGAGGTCAGGAGATCGAGACCATCCTGGCTAACACGGTGAAACCCCGTCTCTACT**
**AAAAAAAAAAAATACAAAAAATTAGCCGGGCGTGGTAGCGGGCGCCTGTAGTCCCAGCT**
**ACTCGGGAGGCTGAGGCAGGGGAATGGCGTGAACCCGGGAGGCGGAGCTTGCAGTGAG**
**CCGAGATCCCGCCACTGCACTCCAGCCTGGGCGACAGAGCGAGACTCCGTCTCAAATCC**
**CCCTACTTGTACAGCTCGTCCATGCCGAGAGTGATCCCGGCGGCGGTCACGAACTCCAG**
**CAGGACCATGTGATCGCGCTTCTCGTTGGGGTCTTTGCTCAGGGCGGACTGGGTGCTCAG**
**GTAGTGGTTGTCGGGCAGCAGCACGGGGCCGTCGCCGATGGGGGTGTTCTGCTGGTAGT**
**GGTCGGCGAGCTGCACGCTGCCGTCCTCGATGTTGTGGCGGATCTTGAAGTTCACCTTGA**
**TGCCGTTCTTCTGCTTGTCGGCCATGATATAGACGTTGTGGCTGTTGTAGTTGTACTCAAG**
**CTTGTGCCCCAGGATGTTGCCGTCCTCCTTGAAGTCGATGCCCTTCAGCTCGATGCGGTTC**
**ACCAGGGTGTCGCCCTCGAACTTCACCTCGGCGCGGGTCTTGTAGTTGCCGTCGTCCTTG**
**AAGAAGATGGTGCGCTCCTGGACGTAGCCTTCGGGCATGGCGGACTTGAAGAAGTCGTG**
**CTGCTTCATGTGGTCGGGGTAGCGGCTGAAGCACTGCACGCCGTAGGTCAGGGTGGTCA**
**CGAGGGTGGGCCAGGGCACGGGCAGCTTGCCGGTGGTGCAGATGAACTTCAGGGTCAG**
**CTTGCCGTAGGTGGCATCGCCCTCGCCCTCGCCGGACACGCTGAACTTGTGGCCGTTTAC**
**GTCGCCGTCCAGCTCGACCAGGATGGGCACCACCCCGGTGAACAGCTCCTCGCCCTTGC**
**TCACCATATTGGCTCGACCTCCAAAAAAGCCTCCTCACTACTTCTGGAATAGCTCAGAGG**
**CCGAGGCGGCCTCGGCCTCTGCATAAATAAAAAAAAAAAAAAAAAAAAAAAAAAAAAAA**
**AAAAATTAGTCAGCCATGGGGCGGAGAATGGGCGGAACTGGGCGGAGTTAGGGGCGGG**
**ATGGGCGGAGTTAGGGGCGGGACTATGGTTGCTGACTAATTGAGATGCATGCTTTGCATA**
**CTTCTGCCTGCTGGGGAGCCTGGGGACTTTCCACACCTGGTTGCTGACTAATTGAGATGC**
**ATGCTTTGCATACTTCTGCCTGCTGGGGAGCCTGGGGACTTTCCACACCCTAACTGACAC**
**ACATTCCACAGCCGTCTC(A)n**

AATTCCCAAGTAACAACAGTGTGTCAGGCACTTGCTAAGGATCCTAAATTGCAGCAAGGCTA
CAATGCTATGGGATTCTCCCAGGGAGGCCAATTTCTGTACGTTCCTTTTGTTATATTGGACC
ACTTATATGGAATTGATTTTTGTTTCTTTTATTCAGATAAATCCACTCCTCCAATTCAAGATTG
TCACCTATCTTGGAAGGGA

#4
GCTCACACAAATTAAAAGCAGCAACTTATAAATAAGTAAGCAAATGAACAAACAGTTCACAAA
AAGAATGAATAAACAGACATACAGAAGTATGTTCAATCTCAGTAGCAATCAGAATGAAGCTT
GGACATAACCTTCTCAGAAAGCCATTTCCAAGTACCTAGGGTCATGAATGGCTTTTATGAAT
CCTGTCTGTATAAACGAGGTGGATGCTAAGGTCAAGGTTAGCGCCATTTTTGGTGCATCCCT
ACAGCATCTTGTATTTGCTGTGTTGTGGCACTTAATCATAATCAGTGGCTTTTGTTGGTTGGT
TGTTCTTCTTGTTTTCTGTCTTCAGCTCTAAACTACAAGCTCTCTGAGGTCAGAGATTATGTC
TTCTGTTATTTTTTCCAAGTCTGCCATCACATCTGGCGTCTGTTTGTTGAATGAATAAATGAA
TAGAGTAAATAGATAGTAATCAAAGGCATAGAAATTAAAGTAA

G**GGCCGGGCGCGGTGGCTCACGCCTGTAATCCCAGCACTTTGGGAGGCCGAGGCGGGC**
**GGATCACGAGGTCAGGAGATCGAGACCATCCTGGCTAACACGGTGAAACCCCGTCTCTA**
**CTAAAAAAAAAATACAAAAAATTAGCCGGGCGTGGTAGCGGGCGCCTGTAGTCCCAGCT**
**ACTCGGGAGGCTGAGGCAGGGGAATGGCGTGAACCCGGGAGGCGGAGCTTGCAGTGAG**
**CCGAGATCCCGCCACTGCACTCCAGCCTGGGCGACAGAGCGAGACTCCGTCTCAAATCC**
**CCCTACTTGTACAGCTCGTCCATGCCGAGAGTGATCCCGGCGGCGGTCACGAACTCCAG**
**CAGGACCATGTGATCGCGCTTCTCGTTGGGGTCTTTGCTCAGGGCGGACTGGGTGCTCAG**
**GTAGTGGTTGTCGGGCAGCAGCACGGGGCCGTCGCCGATGGGGGTGTTCTGCTGGTAGT**
**GGTCGGCGAGCTGCACGCTGCCGTCCTCGATGTTGTGGCGGATCTTGAAGTTCACCTTGA**
**TGCCGTTCTTCTGCTTGTCGGCCATGATATAGACGTTGTGGCTGTTGTAGTTGTACTCAAG**
**CTTGTGCCCCAGGATGTTGCCGTCCTCCTTGAAGTCGATGCCCTTCAGCTCGATGCGGTTC**
**ACCAGGGTGTCGCCCTCGAACTTCACCTCGGCGCGGGTCTTGTAGTTGCCGTCGTCCTTG**
**AAGAAGATGGTGCGCTCCTGGACGTAGCCTTCGGGCATGGCGGACTTGAAGAAGTCGTG**
**CTGCTTCATGTGGTCGGGGTAGCGGCTGAAGCACTGCACGCCGTAGGTCAGGGTGGTCA**
**CGAGGGTGGGCCAGGGCACGGGCAGCTTGCCGGTGGTGCAGATGAACTTCAGGGTCAG**
**CTTGCCGTAGGTGGCATCGCCCTCGCCCTCGCCGGACACGCTGAACTTGTGGCCGTTTAC**
**GTCGCCGTCCAGCTCGACCAGGATGGGCACCACCCCGGTGAACAGCTCCTCGCCCTTGC**
**TCACCATATTGGCTCGACCTCCAAAAAAGCCTCCTCACTACTTCTGGAATAGCTCAGAGG**
**CCGAGGCGGCCTCGGCCTCTGCATAAATAAAAAAAATTAGTCAGCCATGGGGCGGAGAA**
**TGGGCGGAACTGGGCGGAGTTAGGGGCGGGATGGGCGGAGTTAGGGGCGGGACTATGG**
**TTGCTGACTAATTGAGATGCATGCTTTGCATACTTCTGCCTGCTGGGGAGCCTGGGGACTT**
**TCCACACCTGGTTGCTGACTAATTGAGATGCATGCTTTGCATACTTCTGCCTGCTGGGGAG**
**CCTGGGGACTTTCCACACCCTAACTGACACACATTCCACAGCCGTCTC(A)n**

AGAAATTAAAGTAAAATATTTTGGTTTATTTAATAAGGGAAATTAAATTTTAATATATTGTGTGT
TGATAAAAATGTAATGAAGTTAGTATGTTCATTTACTGCTGGTGACA

#5
AGTGTGTGGGTCTAGTGGAGAGATTATCTATCCAAACGCATTTTTGTAATTCCGTACTCTGG
TCCAAAAGTTAATGTAAAAAGCCAATCGAATTGATTAATACTAAAAATGGAACAACTGC

**GGCCGGGCGCGGTGGCTCACGCCTGTAATCCCAGCACTTTGGGAGGCCGAGGCGGGCG**
**GATCACGAGGTCAGGAGATCGAGACCATCCTGGCTAACACGGTGAAACCCCGTCTCTACT**
**AAAAAAAAAATACAAAAAATTAGCCGGGCGTGGTAGCGGGCGCCTGTAGTCCCAGCTAC**
**TCGGGAGGCTGAGGCAGGGGAATGGCGTGAACCCGGGAGGCGGAGCTTGCAGTGAGCC**
**GAGATCCCGCCACTGCACTCCAGCCTGGGCGACAGAGCGAGACTCCGTCTCAAATCCCC**
**CTACTTGTACAGCTCGTCCATGCCGAGAGTGATCCCGGCGGCGGTCACGAACTCCAGCA**
**GGACCATGTGATCGCGCTTCTCGTTGGGGTCTTTGCTCAGGGCGGACTGGGTGCTCAGGT**
**AGTGGTTGTCGGGCAGCAGCACGGGGCCGTCGCCGATGGGGGTGTTCTGCTGGTAGTGG**
**TCGGCGAGCTGCACGCTGCCGTCCTCGATGTTGTGGCGGATCTTGAAGTTCACCTTGATG**
**CCGTTCTTCTGCTTGTCGGCCATGATATAGACGTTGTGGCTGTTGTAGTTGTACTCAAGCT**
**TGTGCCCCAGGATGTTGCCGTCCTCCTTGAAGTCGATGCCCTTCAGCTCGATGCGGTTCA**
**CCAGGGTGTCGCCCTCGAACTTCACCTCGGCGCGGGTCTTGTAGTTGCCGTCGTCCTTGA**
**AGAAGATGGTGCGCTCCTGGACGTAGCCTTCGGGCATGGCGGACTTGAAGAAGTCGTGC**
**TGCTTCATGTGGTCGGGGTAGCGGCTGAAGCACTGCACGCCGTAGGTCAGGGTGGTCAC**
**GAGGGTGGGCCAGGGCACGGGCAGCTTGCCGGTGGTGCAGATGAACTTCAGGGTCAGCT**
**TGCCGTAGGTGGCATCGCCCTCGCCCTCGCCGGACACGCTGAACTTGTGGCCGTTTACGT**
**CGCCGTCCAGCTCGACCAGGATGGGCACCACCCCGGTGAACAGCTCCTCGCCCTTGCTC**
**ACCATATTGGCTCGACCTCCAAAAAAGCCTCCTCACTACTTCTGGAATAGCTCAGAGGCC**
**GAGGCGGCCTCGGCCTCTGCATAAATAAAAAAAATTAGTCAGCCATGGGGCGGAGAATG**
**GGCGGAACTGGGCGGAGTTAGGGGCGGGATGGGCGGAGTTAGGGGCGGGACTATGGTT**
**GCTGACTAATTGAGATGCATGCTTTGCATACTTCTGCCTGCTGGGGAGCCTGGGGACTTTC**
**CACACCTGGTTGCTGACTAATTGAGATGCATGCTTTGCATACTTCTGCCTGCTGGGGAGCC**
**TGGGGACTTTCCACACCCTAACTGACACACATTCCACAGCCGTCTC(A)n**

TAAAAATGGAACAACTGCAAACTAGAACTAATTTGAGAACCCTCCTACTTTCATCCTCAGAGT
TTGTCTAATAAGGACTTTTGGACGACTGAATACATAGCATGTAAGAAAGTGTTAGAAATGGC
TACAAGTCTGATGTGACTGCAACAATATTACTCATAATGTTTTTGAGCACAACAAAAACATAC
AGGGTACATCCACATGCA
